# Mitochondrial dynamics quantitatively revealed by STED nanoscopy with an enhanced squaraine variant probe

**DOI:** 10.1101/646117

**Authors:** Xusan Yang, Zhigang Yang, Ying He, Chunyan Shan, Wei Yan, Zhaoyang Wu, Peiyuan Chai, Junlin Teng, Junle Qu, Peng Xi

**Affiliations:** Department of Biomedical Engineering, College of Engineering, Peking University, Beijing 100871, China; Key Laboratory of Optoelectronic Devices and Systems of the Ministry of Education and Guangdong Province, College of Physics and Optoelectronic Engineering, Shenzhen University, Shenzhen 518060, China; School of Applied and Engineering Physics, Cornell University, Ithaca 14853, United States; School of Life Sciences, Peking University, Beijing 100871, China

**Keywords:** living cell superresolution, squaraine dye, long-term, STED

## Abstract

Mitochondria play a critical role in generating energy to support the entire lifecycle of biological cells, yet it is still unclear how their morphological structures evolve to regulate their functionality. Conventional fluorescence microscopy can only provide ∼300 nm resolution, which is insufficient to visualize mitochondrial cristae. Here, we developed an enhanced squaraine variant dye (MitoESq-635) to study the dynamic structures of mitochondrial cristae in live cells at superresolution. The low saturation intensity and high photostability make it ideal for long-term, high-resolution STED nanoscopy. We demonstrate the time-lapsed imaging of the mitochondrial inner membrane over 50 minutes in living HeLa cells at 35.2 nm resolution for the first time. The forms of the cristae during mitochondrial fusion and fission can be clearly resolved. Our study demonstrates the emerging capability of optical STED nanoscopy to investigate intracellular physiological processes at nanoscale resolution for long periods of time with minimal phototoxicity.

## Introduction

The mitochondrion is the subcellular organelle that provides energy for the cell; thus, it is regarded as “the power plant of the cell” ^1^. Interestingly, the fact that the mitochondrion has its own DNA that differs from that of the host cell may suggest the source of its mysterious origin, in that it may be derived from a kind of ancient bacteria^2^. There is evidence that after its successful incorporation into the host, it altered the evolution of the host cell ^3^ because it could provide it with ATP, which is the only energy form that cells can absorb. Recently, scientists have discovered that cancerous cells can exploit mitochondria to generate more energy for ultra-active metabolism of malignant cells ^4^, while blocking the functioning of mitochondria can stop tumor growth.

The name “mitochondria” comes from the Greek words “mito” (particle) and “chondria” (lines), which describe their various structural characteristics in a cell. Their plasticity and structural dynamics can regulate a number of cancerous cell activities, such as proliferation, migration, and resistance to therapy ^5^. Inside mitochondria, ATP is synthesized at the folds in the inner membranes called cristae. The size of a mitochondrion is usually approximately 1 μm in diameter and 4-10 μm in length, whereas the distance between the cristae in mitochondria is ∼100 nm. Considering the Nyquist-Shannon sampling theorem, one needs at least 50 nm resolution to visualize the gaps between mitochondria cristae. Limited by the diffraction of light, conventional microscopy techniques are insufficient to visualize the structure and dynamics of mitochondria^6^. The mitochondrial intermembrane space can be measured with indirect approaches such as proteomic mapping^7^.

A variety of superresolution techniques have been applied to visualizing submitochondrial structures. For example, Jans et al. discovered the mitochondrial inner membrane organizing system (MINOS) with STED^8^. With 3D-STED, Schmidt et al. imaged the mitochondrial cristae at isotropic resolution^9^. Using 3D stochastic optical reconstruction microscopy (STORM), Huang et al. showed the 3D ultrastructure of the mitochondrial network in a fixed cell^10^. Furthermore, to resolve the dynamics of mitochondria in living cells, Shim et al. developed lipophilic cyanine dyes ^11^, while Tang et al. developed photoactivatable Znsalen complex^12^. Ishigaki et al. reported the rich dynamics of mitochondria with a Rhodamine derivative^13^. However, because they are limited by the spatial resolution, the cristae can only be visualized as lamellar curtain-like structures. In both cases, the spatial resolution was insufficient to resolve the individual cristae for further analysis. Dannie et al. also demonstrated label-free third-order photoacoustic (PA) nanoscopy of the cytochromes in the inner mitochondrial membrane in fibroblasts at 88 nm resolution^14^. Recently, we have reported the use of Hessian structured illumination microscopy (SIM) to investigate mitochondria dynamics, in which the cristae can be clearly visualized ^15^. Even though it has a fast imaging speed, the highest spatial resolution for SIM (∼90 nm) is still not enough to measure the sizes and distances in cristae during their evolution^16^. The high spatial resolution (∼ 50 nm) and temporal resolution (∼ 1 frame per second) make STED the most promising choice for the study of mitochondria, which is a tiny cell inside another cell^8,9,17,18^.

There are two constraints prohibiting the application of STED to long-term live cell imaging. (1) cells have a limited tolerance for light exposure due to phototoxicity. Mitochondria are more sensitive to light than other cellular organelles due to the existence of photoreceptors encoded by the pro-fusion gene optic atrophy 1 (OPA1); excessive light exposure can cause mitochondrial dysfunction and mitophagy^19,20^. (2) For STED nanoscopy, high intensity light is required to achieve improved resolution because the fluorescence in the “donut” area must be converted to stimulated emission (laser process) through high power laser illumination ^21^. Moreover, the existing dyes for labeling mitochondria (such as MitoTracker) have several disadvantages, as they are not photostable enough to endure long-term STED imaging and the efficiency is generally too low to generate effective STED processing^13,22^. With the recent developed SNAP substrates and benzylguanine derivative tags, Bottanelli et al. have imaged the mitochondria and ER network with two-color STED for 36 seconds^23^. Very recently, Jakobs group has reported a new cell line expressing mitochondrial protein fused to SNAP-tag, to enable high resolution STED of mitochondrial cristae in live cell for 2 minutes (every 15 second per frame, 8 frames) ^24^.

To address these challenges, in this work, we have developed a new squaraine dye derivative (MitoESq-635) that is compatible with live cells and produces low phototoxicity. The primary advantage of this dye is that it can be easily depleted by a STED laser at relatively low power, which allows the cells to stay in their native state during STED imaging. With this ESq-STED dye, we achieved a spatial resolution of 35.2 nm during time-lapse imaging of mitochondrial cristae dynamics for the first time. The fusion and fission processes can be clearly visualized, and the evolution of the cristae structure can be observed over time, which is impossible with other current imaging techniques. Because it benefited from the photostability of MitoESq-635, STED imaging of a 3D stack revealed its axial distribution. Because of the capability of imaging mitochondria cristae dynamically at superresolution, we believe the MitoEsq-635 dye will be widely applied for the study of mitochondria-related cell behaviors and the discovery of the origin of mitochondria.

## Results

### Imaging live cells with the modified squaraine dye

We have recently developed an enhanced squaraine variant dye (MitoESq-635) for the labeling of mitochondria in live cells. A hexylamidophenylarsenicate chain is conjugated to the sulfide atom at the central position of the four-membrane ring in the squaraine dye (see Supplementary Note 1), which can be used as a protein label for the plasma membrane. Due to the fast binding of phenylarsenicate to vicinal dithiols and the prioritized targeting of mitochondria by the dye molecules, incubating live cells with MitoESq-635 for a few minutes is enough to label the plasma membrane with high density. Vicinal-dithiol-containing proteins (VDPs) containing two active thiol groups in the vicinity can be covalently labeled by a phenylarsenicate conjugate ^25,26^. To verify that the enhanced squaraine variant VDP probe (MitoESq-635 in Figure 1) is specifically bound to the membrane proteins in the mitochondria, HeLa cells treated with MitoESq-635 for 5 minutes were imaged with a confocal laser scanning microscope. The fluorescence signals were concentrated in the eccentric perinuclear regions, whereas little signal was present in the cytoplasmic and nucleus regions. Imaging of the colocalization of MitoESq-635 and MitoTracker in HeLa cells suggested that the probe was mainly concentrated in mitochondria (see Supplementary Figure 1). During colocalization studies with ER and lysosome trackers, some localization within the ER was also observed^27^. Additionally, MitoESq-635 is widely compatible with different cell lines. We tested and found that it is compatible with cultured HeLa, MCF7, RAW 264.7, and U2OS cell lines as well as primary neuron cells (see Supplementary Figure 2). The covalent binding of MitoESq-635 with mitochondrial VDPs was further verified by SDS-PAGE, fixed cell washing and colocalization experiments (see Supplementary Figures 3 and 4).

**Figure 1.**
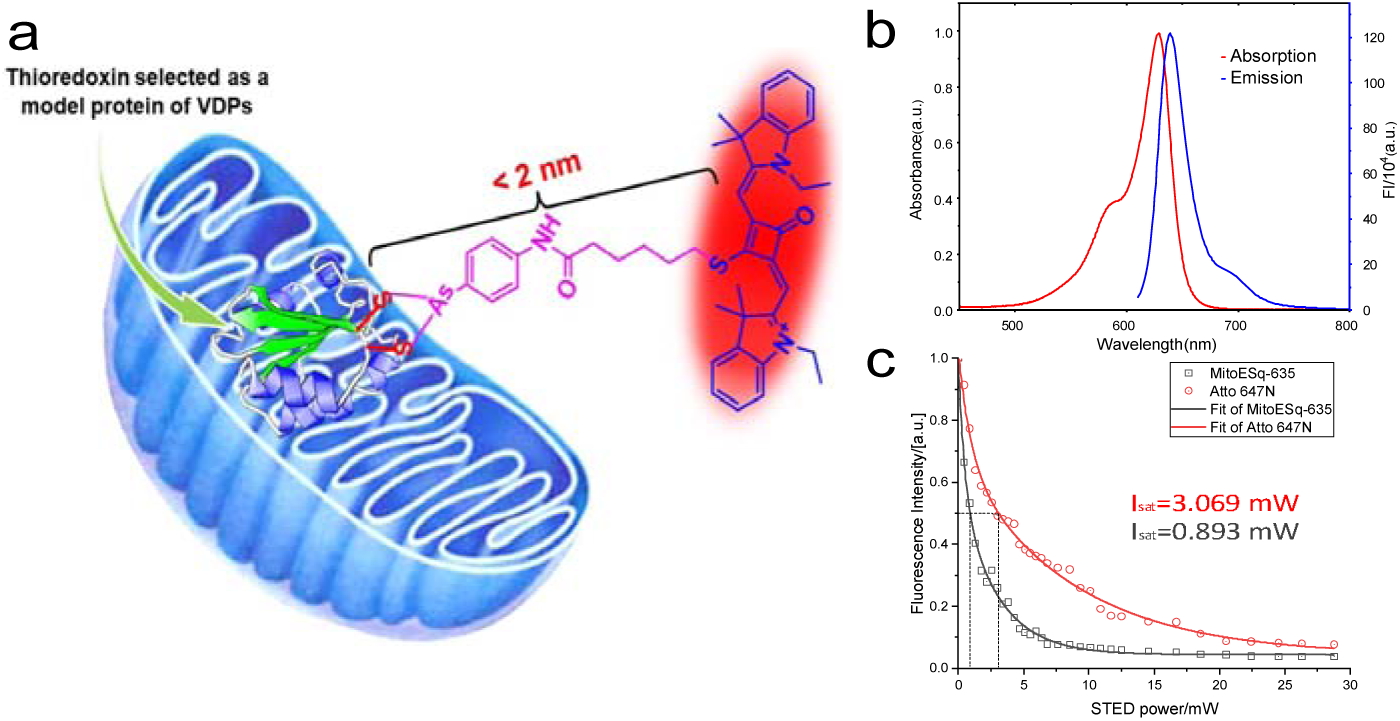
Development of a highly photostable and minimally phototoxic bright cell-permeable enhanced squaraine probe for mitochondria inner membrane labeling. (a) Chemical structure of MitoESq-635 used for the specific labeling of VDPs. (b) Absorption and emission spectrum of the enhanced squaraine dye, for which a 775 nm pulse laser can be employed for depletion and a 635 nm laser for excitation during STED setup. (c) The detected fluorescence signal as a function of the depletion beam intensity; the excitation beam was 635 nm with a 5 ps pulse width and 80 MHz, and the STED beam was 775 nm with 600 ps pulse width and 80 MHz.

**Figure 2.**
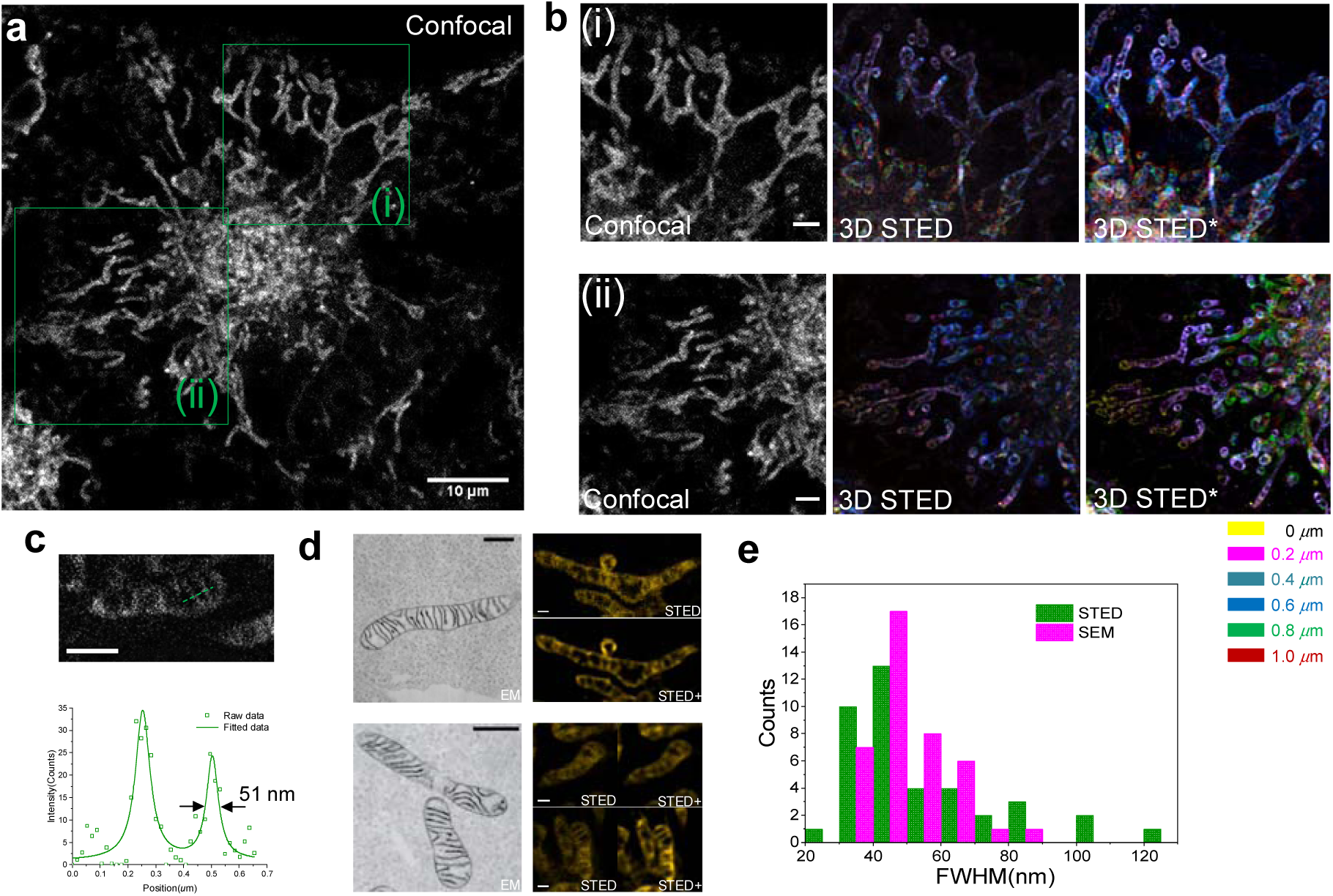
3D superresolution imaging of the mitochondria in HeLa cells with MitoESq-635. (a) 2D confocal image of mitochondria labeled with an enhanced squaraine probe in living HeLa cells. (b) The color-coded STED imaging of the 3D stack (raw data) of the corresponding green boxed area (i, ii) in (a); STED* is the corresponding deconvoluted image of the STED image. (c) Resolution in mitochondrial cristae. (d) In the STED images and the corresponding deconvoluted images (STED+), the cristae structures in living cells can definitely be resolved and can be compared to the previously reported APEX 2 labeled EM images that were reprinted with permission from ref. ^30^; scale bars in EM and STED images are 500 nm and 1 μm, respectively. (e) Cristae width distribution histogram based on STED and SEM images. The raw data for the STED images (1024×1024 pixels) in different layers of (b) are shown in **Supplementary Videos 1 and 2,** in which the z-step size is 200 nm, 635 nm is the excitation wavelength, and the power of the 775 nm laser used for depletion at the back focal plane of the objective was 5.2 μW and 36 mW, respectively; the pixel dwell time was 4 μs. Scale bars in (a), (b) and (c) are 10 μm, 2 μm and 1 μm, respectively. The scale bars of the EM and STED fluorescence images in d are 500 nm and 1 μm.

**Figure 3.**
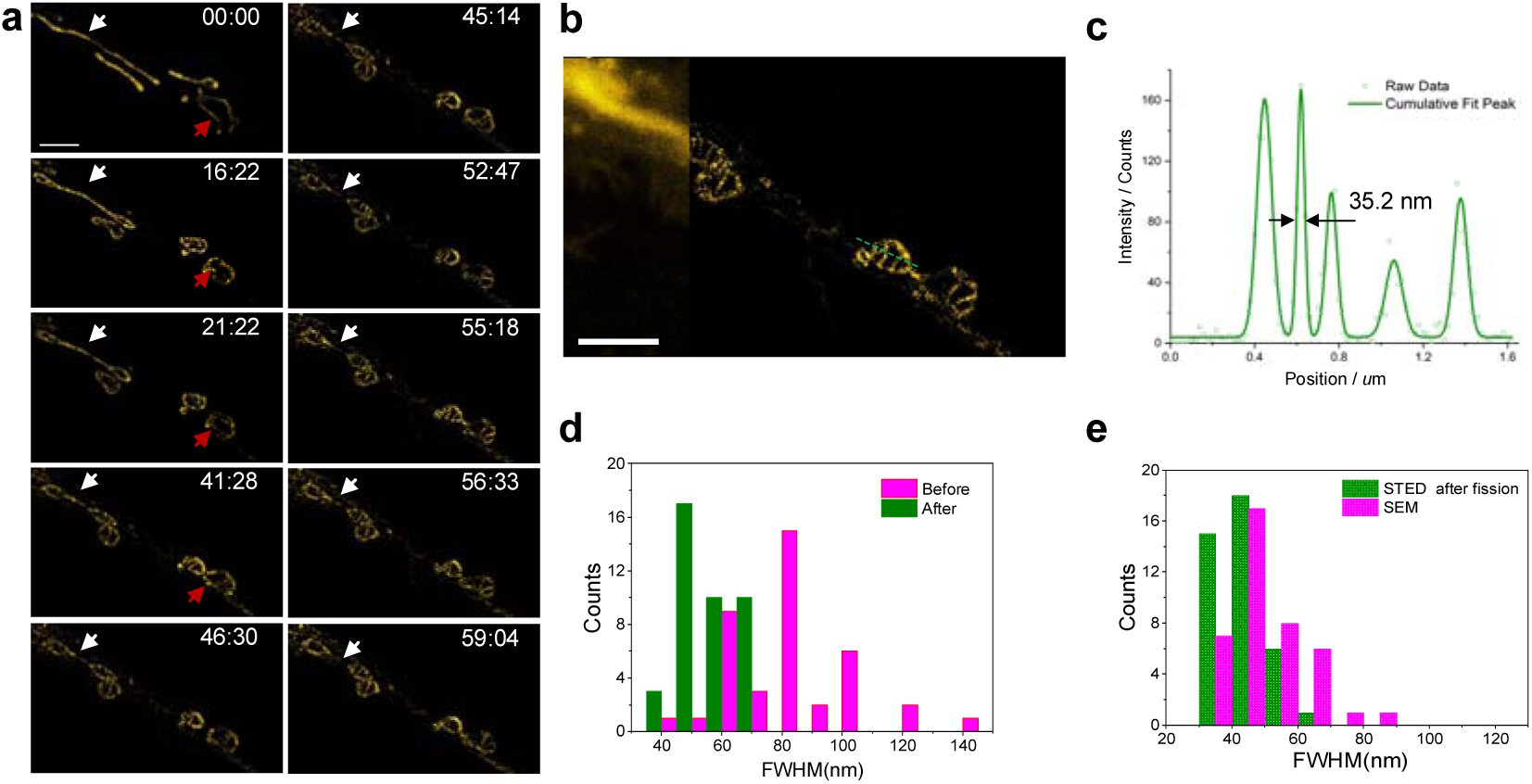
Confocal (a, left) and STED (a, right) imaging of the mitochondrial cristae. (a) Time-lapsed STED imaging of the mitochondria. The white and yellow arrows show typical mitochondrial fission processes. (b) Comparison of confocal (left) and STED (right) imaging. The intensity profile across the green dashed line in (b) is shown in (c). A 35.2 nm resolution can be obtained with STED superresolution microscopy. (d) Distribution of the widths of the cristae. Green bars: connective tubule width distribution histogram after fission. Pink bars: connective tubule width distribution histogram before fission. (e) Cristae width distribution histogram after fission in STED and EM. Scale bar, 2 μm.

**Figure 4.**
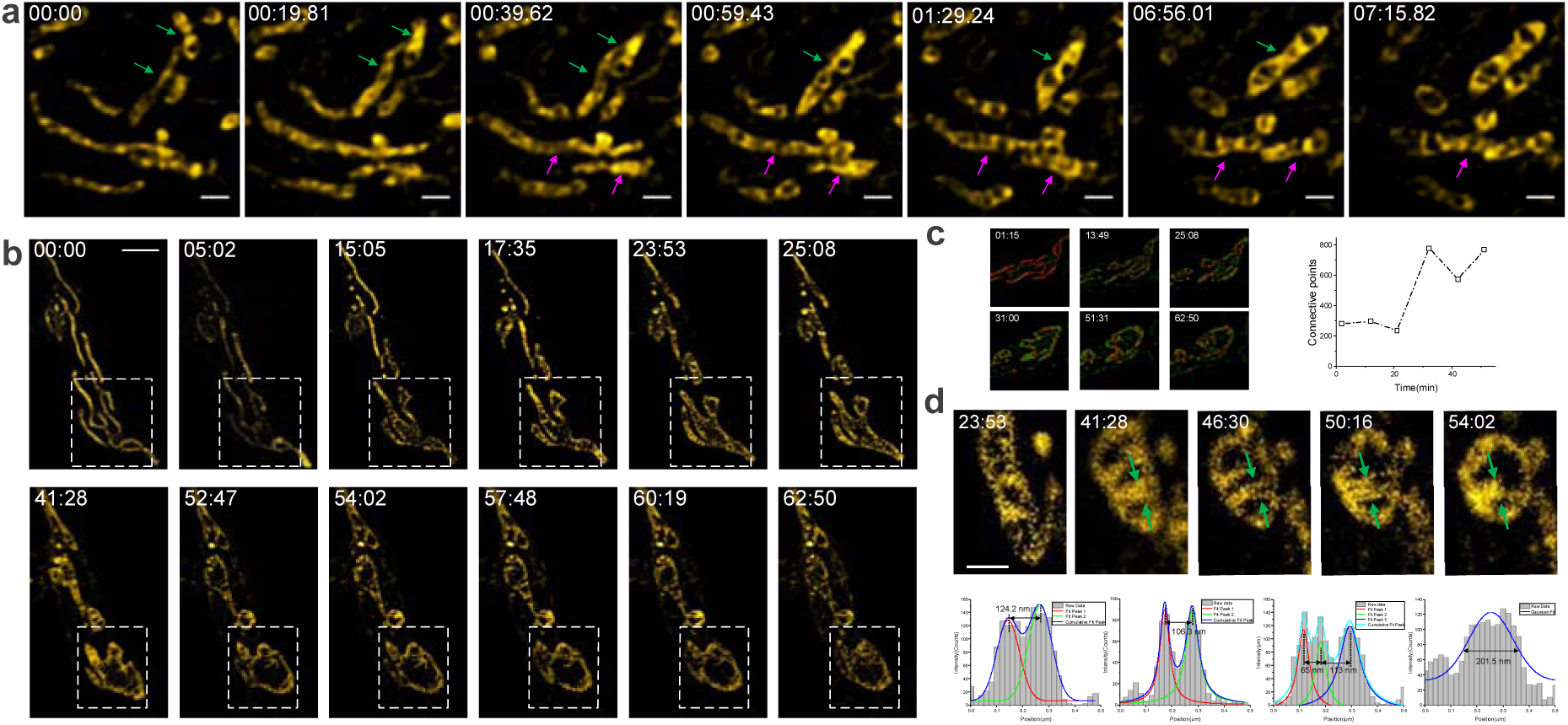
Mitochondrial fission and fusion processes observed with STED optical nanoscopy. (a) The mitochondrial fission process revealed cristae dynamics (periodic in the matrix) during fusion (the raw data is shown in Supplementary Video 5). (b) Skeletal image of mitochondria quantitatively revealing cristae dynamics during fusion (the raw data can be seen in Supplementary Video 6). Scale bar= 5 μm. (c) Skeletal image of mitochondria quantitatively revealing cristae dynamics during fusion. (d) Cristae dynamics during conversion from a line shape to a bubble. The cross-section plots represented by the green arrows are shown in the lower row. Scale bars in a, b and d are 1 μm, 2 μm and 1 μm, respectively.

### Photostability and stimulated emission saturation intensity of the dye

Compared with standard gold live cell mitochondrial probes, MitoESq-635 exhibits much more robust photostability when exposed to a focused laser in confocal microscopy (MitoTracker Green, see Supplementary Figure 5 left) and in STED nanoscopy (MitoTracker DeepRed, see Supplementary Figure 5 right). ATTO 647N is the standard fluorescent label in the red spectral region commonly used for STED nanoscopy because of its strong absorption, high photostability, and high resolution at relatively low STED power. However, the squaraine-STED dye has a lower saturation intensity at 0.893 mW, which is ∼3.4-fold higher than that of Atto 647N (Figure 1). The saturation intensity of the organic fluorophore (Atto 647N) was measured to be P = 3.069 mW when light reduced the fluorescence by half. This is comparable with values already published (P = 9 mW for CW STED^28,29^). Because of its low saturation intensity, the MitoESq-635 probe exhibited no marked toxicity in HeLa cells at a concentration of 1 μM after 1 hour of incubation during STED superresolution imaging microscopy (see Supplementary Figure 6). The low saturation intensity, extended photostability, and low toxicity make the dye very suitable for long-term STED imaging in live cells.

Next, the probe was applied to 3D stack STED imaging in live cells. Previously, it was challenging to perform STED in 3D stacks because of photobleaching during imaging. As shown in Figure 2, the regions (i, ii) in panel a were scanned with a z-step of 200 nm, and 3D stack STED imaging was obtained by the construction of different layers of STED images represented by different colors. The raw data were used to obtain fine STED imaging of mitochondria, which could be optimized through deconvolution. From the magnified STED image of a single mitochondrion, it can be observed that the probe was mainly localized in the cristae, which were formed by the folding of the inner membrane of mitochondria. The morphological structures in cristae were resolved by STED nanoscopy and matched well with those in previously reported APEX 2-labeled EM images (panel c, d)^30^, and their widths were as small as ∼51 nm (panel c), which was in line with that observed in the EM images (panel e).

### Subcellular dynamic nanoscopic imaging with MitoEsq-635

While detailed structural information for intracellular membranes at nanoscales has emerged from electron microcopy^31-33^ as well as localization superresolution microscopy^34,20^, it is still unclear how different forms of mitochondria have evolved. A fast superresolution technique with sufficient spatial and temporal resolution is highly desired. To study the highly dynamic and subtle morphological changes within mitochondria, we developed a long-term STED nanoscopic imaging strategy to visualize mitochondrial membranous dynamical structures in living cells. Taking advantage of the live cell compatibility of MitoESq-635, the rapid nanoscale spatiotemporal mitochondrial dynamics in a living HeLa cells was captured with STED (1.2 s per frame, 700 × 700 pixels, 12.6 μm ×12.6 μm, STED beam of 33.6 mW at 775 nm before the objective) long-term (over 10 minutes) (**see Supplementary Video 3**), during which the mitochondria showed healthy behavior with rare changes to spheroidicity. Additionally, we were able to record the longer term (over 60 minutes, 3.9 s per frame with 71.5 s of dark recovery) nanoscale spatiotemporal dynamics of the mitochondrial inner membrane with STED nanoscopy at a resolution of 35.2 nm (for the raw data, see **Supplementary Video 4**) to observe mitochondrial fission. As shown with the white arrow, the thin and elongated mitochondria first form bubble structures at 16:22; then, the cristae grow quickly inside these bubbles, and they separate to form individual small mitochondria with rich and constantly changing internal cristae structures (21:22-56:33) (Figure 3).

The time-lapsed STED images revealed thin, extended tubular intermediates connecting neighboring mitochondria both before fission and after fission (Figure 3). The average widths of these tubular structures before and after fission were approximately 85 and 42 nm, respectively. Additionally, the cristae width after fission was measured to be 43 nm, which agrees well with the reported EM results^35^. Remarkably, a high power (∼1 mW) excitation beam is needed to obtain adequate fluorescence signals from MitoTracker DeepRed, which will cause a marked change in the intracellular condition (mitochondrial shape change from a network to spheroidicity within 2 minutes, see Supplementary Figure 7) because the wavelength is at the absorption peak maximum of the probe. The quenching beam (STED) could also cause intracellular condition changes at a light dose to a certain extent. If the power of the quenching beam is further increased, the fluorescence molecule could melt under the high depletion beam power of 51 mW (see Supplementary Figure 8). With MitoESq-635, rare photobleaching and mitochondrial shape variations are observed upon exposure to STED scanning for over 10 minutes (1.2 s per frame, 700 × 700 pixels, 12.6 µm × 12.6 µm, STED beam of 30.2 mW at 775 nm), as shown in Supplementary Figure 5. In contrast, the fluorescence signal from MitoTracker Green dropped very quickly due to significant photobleaching (>70% in 300 scans, as shown in the third image in Supplementary Figure 5), making it unsuitable for long-term live cell STED imaging.

We also observed the fusion process of mitochondria, as shown in Figure 4 (also see Supplementary Videos 5 and 6). As highlighted by the white dashed boxed area, the mitochondria were initially in a typical line structure. Then, the cristae begin to form and grow rapidly (15:05) in association with merging and combining with a larger area (23:53-41:28). The merge makes the finger-like structures disappear, and a hollow structure with several shoots containing cristae forms (52:47-60:19). The shoots then form a complicated network of cristae (44:00). The arrows point to several areas where the rod-like isolated mitochondria become expanded with newly formed cristae and fuse into one large mitochondrion with several bubbles. After mitochondrial fusion, the cristae membrane begins to remodel. All of these changes are in line with those seen in the EM images reported previously, as shown in Supplementary Figure 9. The inner mitochondrial membrane is compartmentalized into numerous cristae, which expand the surface area of the inner mitochondrial membrane to enhance its ability to produce ATP. The oxidative phosphorylation system (OXPHOS) facilitates energy conversion for ATP production so that the mitochondria can supply energy for other subcellular organelles, which make mitochondria the indispensable ‘power plants’ of eukaryotic cells. Unlike the fission process shown in Figures 3 and 4, in which the mitochondria were ultimately separated, in these figures both fusion and fission processes occur simultaneously.

## Conclusion

Mitochondria play important roles in cells because they generate energy for a variety of cellular and developmental processes. Mitochondrial dysfunction is associated with numerous severe diseases, including several devastating neurodegenerative diseases. Furthermore, the degree, functional relevance, and molecular causes of the heterogeneity of mitochondrial structure and function in healthy and stressed single cells requires both molecular/biochemical tools and advanced superresolution living cell microscopy. When responding to different cellular statuses, mitochondria constantly change the morphologies of their outer and inner membranes to regulate energy production. To investigate their dynamic behavior in live cells, fast and high-resolution superresolution microscopy techniques are required. Because the distance between mitochondrial cristae is less than 90 nm, SIM cannot fully resolve the cristae due to limited resolution enhancement. Single-molecule localization microscopy techniques can provide 20-50 nm spatial resolution, but their poor temporal resolution can lead to significant motion blur during the imaging process. Although STED can provide high spatial and temporal resolution at the same time, the current STED dyes for mitochondria are generally incompatible with live cells during long-term STED imaging due to high light intensity-induced phototoxicity and photobleaching. In this work, we report the use of MitoESq-635, which is a novel squaraine dye specifically designed for long-term live cell STED imaging of mitochondria. MitoESq-635 probes have superior photophysical properties compared organic dye ATTO 647N probes, which are commonly employed in STED nanoscopic imaging. Its colocalization with MitoTracker demonstrated that MitoESq-635 can label the membrane of mitochondria. Here, we achieved 35 nm spatial superresolution at a STED imaging speed of ∼1 fps. The dynamic imaging of live cells over 50 minutes clearly revealed the fusion and fission processes of mitochondria. Because of the low saturation intensity, STED imaging of 3D stacks can reveal the ultrastructure of mitochondria in live cells. Overall, its labeling specificity and performance in low power STED make MitoESq-635 highly attractive as a next-generation standard for long-term superresolution imaging of mitochondria.

## Acknowledgments

This work was supported by the National Key Research and Development Program of China (2017YFC0110202), the National Natural Science Foundation of China (61475010, 61525503, 61729501, 61875131, 6187030647, 61620106016/61835009), the Beijing Natural Science Foundation (JQ18019), the (Key) Project of the Department of Education of Guangdong Province (2015KGJHZ002/2016KCXTD007), the Guangdong Natural Science Foundation Innovation Team (2014A030312008), and the Shenzhen Basic Research Project (JCYJ20170818100931714/ JCYJ20150930104948169/JCYJ20160328144746940/JCYJ20170412105003520). The authors would like to thank Dr. Kuangshi Antony Chen, Dr. Luke Lavis and Dr. Xu Zhang for helpful discussions regarding this work.

## Author contributions

X.Y., Z.Y., J.Q. and P. X. initiated the project. Z.Y. developed the enhanced squaraine probe. X.Y., Z.Y., and P. X. designed the experiments. X.Y., C.S., P. C. and Y. H. prepared living cells labeled with the enhanced squaraine probe and performed STED superresolution experiments in living cells. X.Y. and W.Y. constructed the microscope setup and designed the optical system for the dye property measurements. J.Q. and P.X. supervised the project. X.Y., Z.Y. and P.X. analyzed the data. X.Y., Z.Y. and P. X. wrote the manuscript with contributions from all authors.

## Methods and materials

### UV/Vis absorption and fluorescence spectroscopy measurements

A stock solution of MitoESq-635 (1 mM) was prepared in DMSO solvent and then diluted with ethanol to 1 μM. The UV/Vis absorption spectra of MitoESq-635 were measured using a Perkin Elmer 3000 spectrophotometer (PE, USA), and the fluorescence emission was measured with Horiba spectrofluorometric equipment (Horiba) with a Xenon lamp, a regular PMT detector and an In/Ga/As detector for the NIR II measurements.

### Preparation of reduced BSA (rBSA) and oxidized BSA (oBSA)

rBSA was prepared by reducing BSA solution with DTT (1.0 mM) overnight at 4 °C, and oBSA was made by oxidizing BSA with H_2_O_2_ (100 μM) at 25 °C for 10 minutes. These samples were then diluted 50 times with distilled water, and the proteins were recovered by precipitation in 50% acetone at −20 °C for 2 hours.

### Cell culture

The cell lines, including HeLa cells and Raw 264.7 cells, were cultured at a suitable density (moderate) in DMEM media. All culture media were supplemented with penicillin (100 units/mL), 10% (v/v) FBS (WelGene) and streptomycin (100 mg/mL), and cells were incubated in a 5% CO_2_ atmosphere at 95% humidity and 37 °C. The cells were placed in confocal dishes 24 hours prior to the experiments.

### Confocal microscopy

The selected cell lines were seeded on cover slips or glass-bottomed dishes (SPL Lifesciences Co., Ltd.) before they were grown to a suitable density (24 hours) via incubation in a humid atmosphere containing 5% (v/v) CO_2_ at 37 °C. The cells were coincubated with MitoESq-635 or mitochondrial, lysosomal, or ER trackers for different times (e.g., 30 minutes) in a humid atmosphere with 5% (v/v) CO_2_ at 37 °C. Then, the cell images were obtained using a confocal laser scanning microscope (Leica SP2 and SP8, Leica, Germany). Related information is available in the figure captions.

### SDS-PAGE and fluorescent imaging of gels

The selectivity of MitoESQ-635 for proteins and cells was identified by 10% SDS-PAGE experiments. The different protein samples were treated with DTT or H_2_O_2_ and were incubated with MitoESQ-635 in PBS buffer at 37 °C for 1 hour. After labeling, the obtained samples were precipitated in 50% (v/v) acetone for 2 hours at −20 °C and then mixed with SDS-PAGE loading buffer containing tris (2-carboxyethyl) phosphine (TCEP), and electrophoresis was carried out immediately. The gel was observed and imaged using a fluorescent scanner (Tanon-5220S, Shanghai, China) with green light excitation with a band path filter with a range from 635 to 675 nm.

### STED superresolution of live cells

STED imaging was performed with a Leica TCS SP8 STED 3X system equipped with a white light laser for excitation and a 775 nm pulsed laser for STED depletion. A 100x oil-immersion objective (Leica, N.A. 1.4) was employed. Each frame was obtained at 1 s, and the image series were obtained at a 1-minute interval.

