## Supplemental Figures for "Mitochondrial dynamics quantitatively revealed by STED nanoscopy with an enhanced squaraine variant probe"

### Supplementary Figures

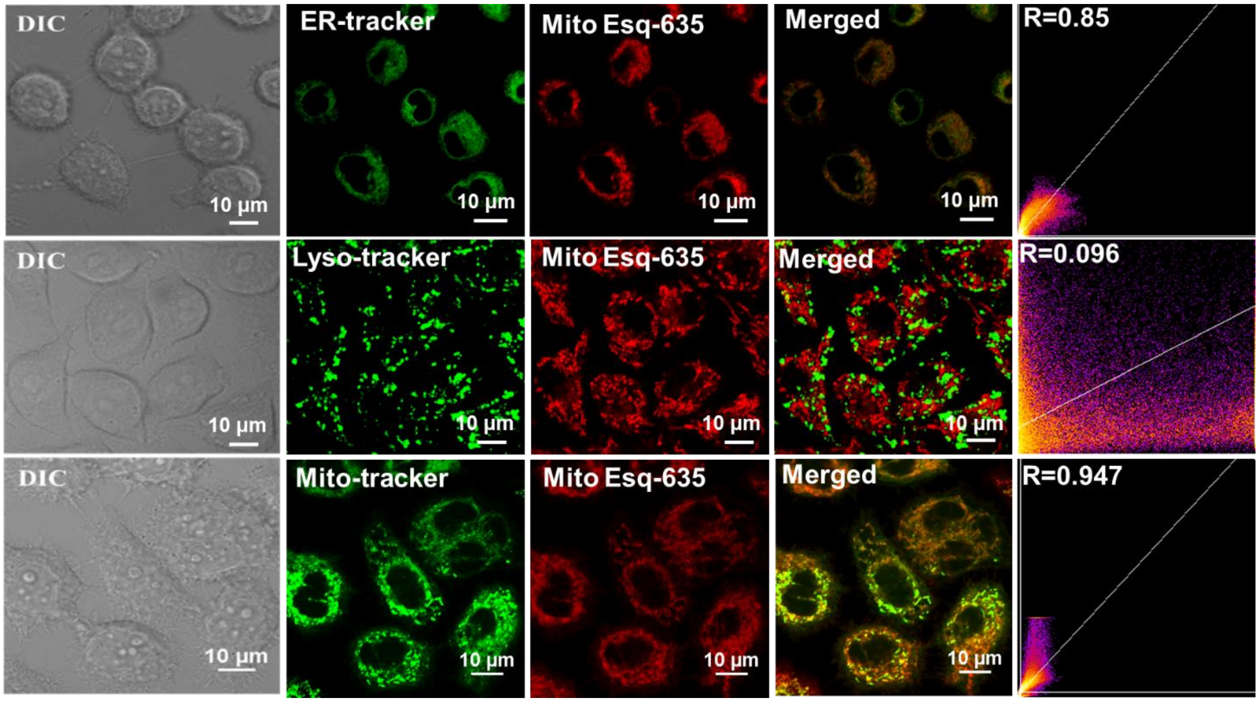


**Supplementary Figure 1**

Co-localization experiment employing MitoTracker Green (Rhodamine 123) as golden standard for mitochondrial marker. The first column, DIC image; the second column, ER- Lyso- and Mito-Tracker green labeled HeLa cells; c, g the third column, enhanced squaraine dye labeled HeLa cells; the fourth column, merged images; the fifth column, Pearson's coefficients, excitation wavelength: Rhodamine 123, Lyso- and ER- trackers (488 nm), MitoESq-635 (633 -nm), detect range of Rhodamine 123 Lyso- and ER-trackers (500-560 nm); MitoESq-635 (640-800 nm), Scale bar, 10 μm.

From Supplementary Figure 1-3, it can be demonstrated that MitoESq-635 can preferentially target for mitochondria in live cells. HeLa cells were subjected to incubating with Mito-, Lyso-, ER- trackers and MitoESq-635, respectively. From the confocal imaging results obtained by a Leica SP8 confocal Laser scanning microscope, the probe revealed the best overlapping imaging with Mitotracker (Rho123).

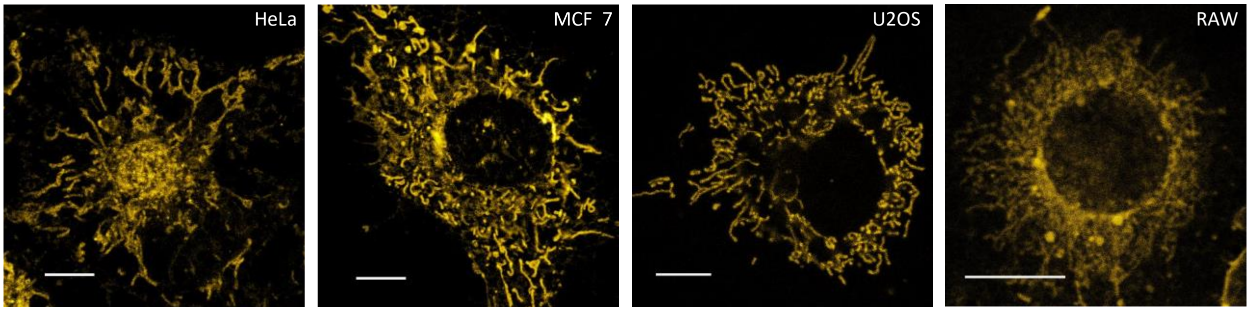


#### Supplementary Figure 2

MitoESq-635 labelling in HeLa, MCF7, Raw264.7, and U2OS cell lines. Scale bar, 10 μm.

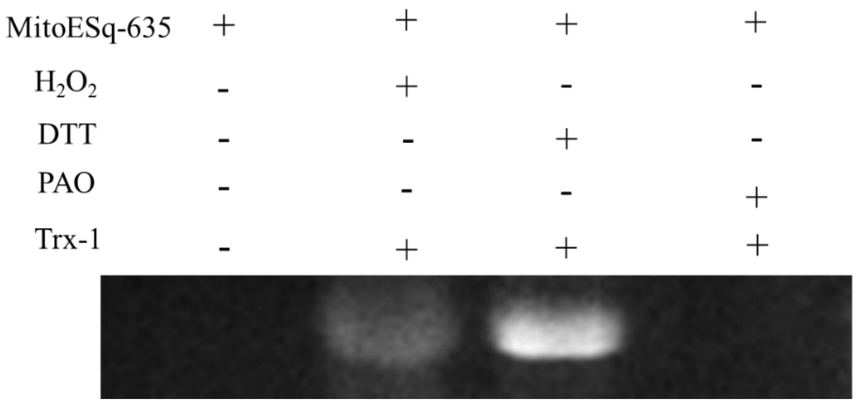


#### Supplementary Figure 3

Identification of the covalent binding of MitoESq-635 to Thioredoxin (Trx-1) proteins under different conditions; $H_2O_2$ (10 mM), DTT (10 mM), PAO (10 mM) were added into Trx-1 protein (20 μM) solution separately, and incubated for 3 hours at 4 ℃. After incubation, each sample was labeled with probe (50 μM) at 37 ℃ for 40 minutes. From the SDS-PAGE results, it can be seen that MitoESq-635 only in the presence of reduced Trx-1 prepared by DTT, the probe can be covalently bound to the vicinal dithiols in the protein. In the presence of PAO, the vicinal dithols on the Trx-1 proteins were masked by PAO to inhibit the reactions of Trx-1 with MitoESq-635. When $H_2O_2$ was added into above solution, the vicinal dithiols were oxidized into disulfide residue in the protein which disables the labeling reaction of MitoESq-635 with Trx-1. Therefore, DTT reduced the Trx-1 to provide rTrx-1 (with vicinal dithiols) to promote the labeling of MitoESq-635 to Trx-1.

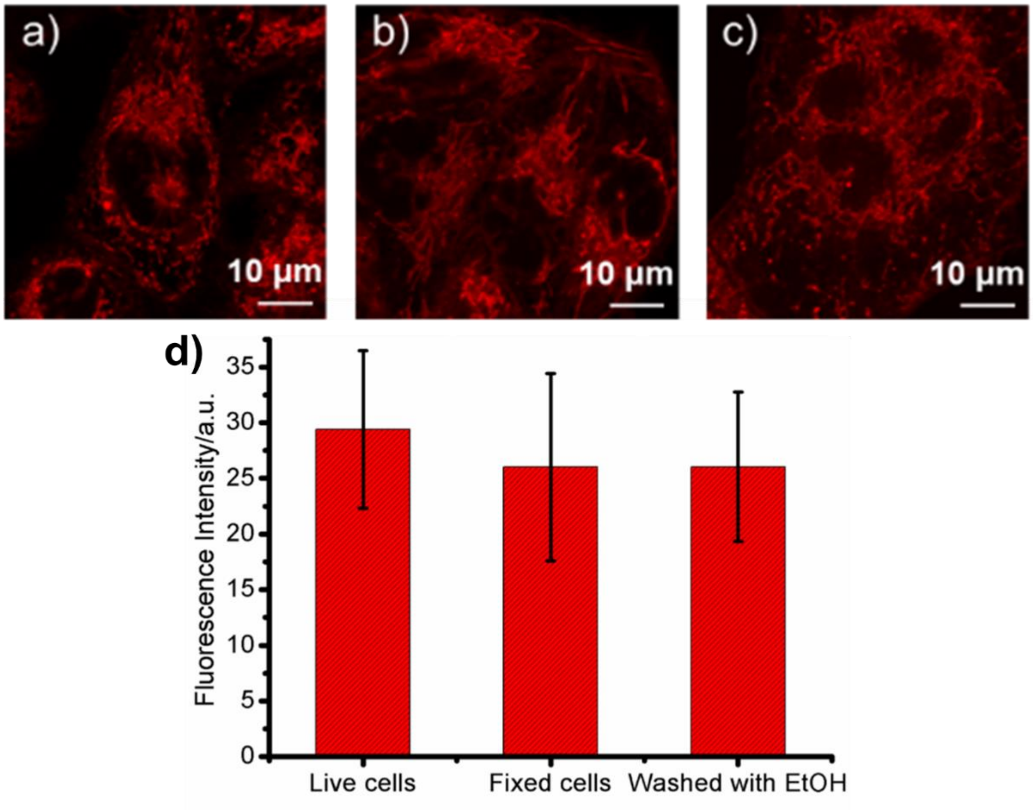


#### Supplementary Figure 4

Proof of covalent binding of **MitoESq**-635 to the VDPs inside cells; confocal images of Hela cells stained with MitoESq-635 (0.5 μM) before (live cells) (a) and after cell fixation, (fixed cells) (b), after fixation, cells were incubated with pure ethanol (EtOH) for 5 minutes (washed out the free probe), then rinsed by PBS for three times (c); and (d), quantitative measurement of fluorescence intensity in (a), (b) and (c). Compared with live HeLa cells, the probe was mainly localized in mitochondria with rare dispersion in the fixed HeLa cells, and even washed with pure ethanol to remove the free probe molecules. It is demonstrated that the probe molecules linked with the VDPs in mitochondria through phenarsenicate moiety very steadily.

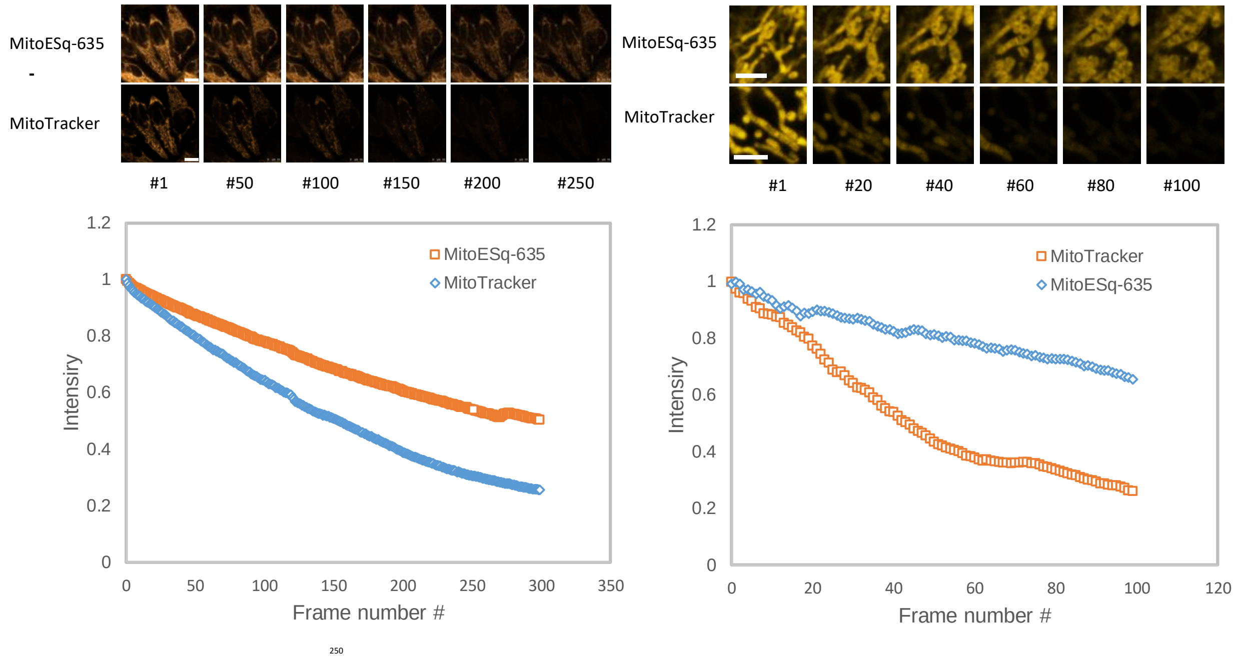


**Supplementary Figure 5.**

Left: Fluorescent imaging changes of HeLa cells co-stained by MitoTracker (Rhodamine123) (upper row, excited at 488 nm, laser power 1.97 μW, collected at the range of 500-560 nm) and MitoESq-635 (lower row, excited at 633 nm, laser power: 1.97 μW, collected at the range of 645-680 nm) with different laser scan time periods under a confocal laser microscope (Commercial Leica SP8 system), image acquisition speed of 1frame per second. Scale bar,10 μm.
Right: The photon bleaching curving of 1.0 μM MitoESq-635 and 0.1 μM MitoTracker DeepRed. The fluorescence intensity of MitoESq-635 and MitoTracker DeepRed was measured by a STED microscopy (TCS SP8 STED 3X). The excitation is 640 nm with the power of 1.1 μW, STED is 775 nm with the power of 36 mW. The detection is HyD3 (647-750 nm, gain 80, no gating). Scale bar,10 μm.

From above results, both MitoTracker (Rho123) and MitoESq-635 were initially concentrated in mitochondria; when the laser was applied to scan the cells for different time, it can be seen that green fluorescence of MitoTracker (Rho123) gradually diffused through the whole cells with reduced fluorescence intensities, while MitoESq-635 was still localized in the mitochondrial region with rare fluorescence decrease. It can be proposed that with the continuously scanning of laser (2.9 μW), MitoESq-635 still localized stably at the mitochondria area due to covalently binding with VDPs, while Rho123 diffused markedly out of mitochondria ascribing to be dependent on membrane potential of mitochondria.

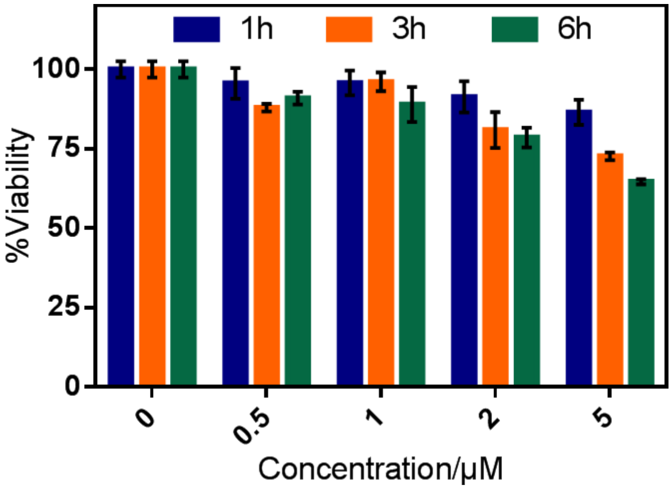


**Supplementary Figure 6.**

Viability experiment on HeLa cells when incubating with MitoESq-635.

HeLa Cells were cultured with the probe with different concentrations (0, 0.5, 1.0, 2.0, 5.0 μM) for different time periods (1, 3 and 6 hours). It can be found in the figure that the probe exhibited low toxicity to HeLa cells, because cell viability was more than 70%, even at the concentration of 5.0 μM for 6 hours' incubation. For the present experiment, low concentration (1.0 μM) with 1 hour's incubation were employed to prepare for STED super-resolution imaging microscopy, which demonstrated that our probe showed no marked toxicity to HeLa cells.

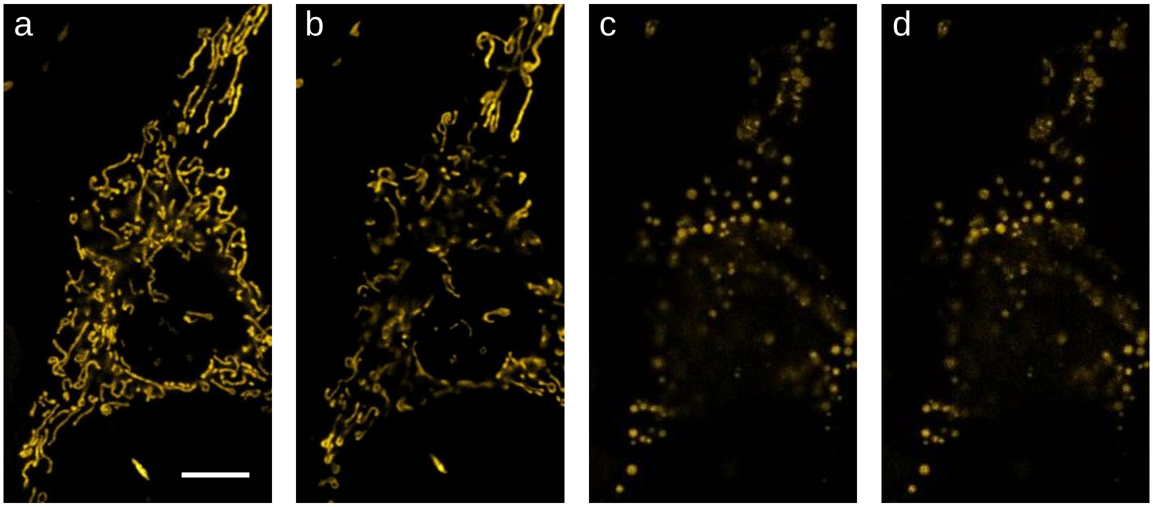


**Supplementary Figure 7.**
Laser photo-toxicity induced cell death by high power dose excitation (1.0 mW) in confocal microscopy. (a-d) Scale bar, 10 μm.

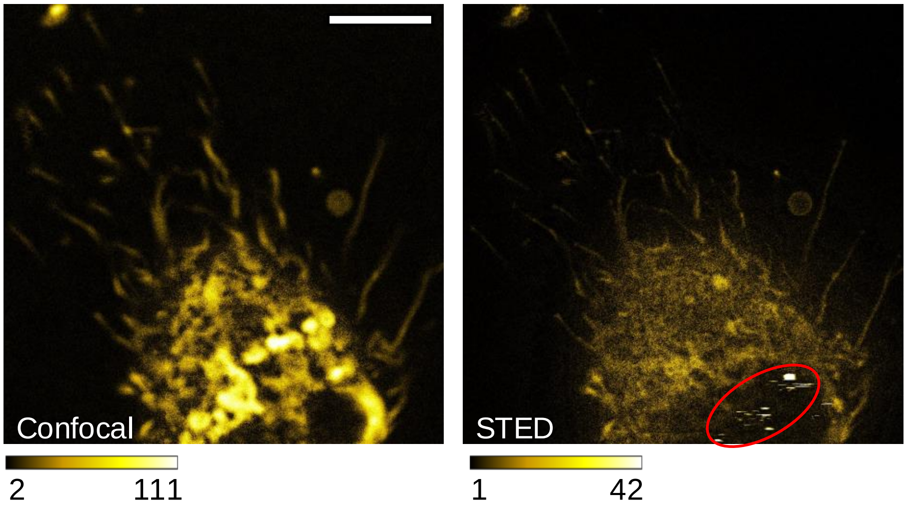


**Supplementary Figure 8.** Melting caused by high dose depletion beam power in STED microscopy. Scale bar, 5*u*m. Depletion beam power, 51mW. The molecules in red circle of STED image is melting induced by high dose depletion beam.

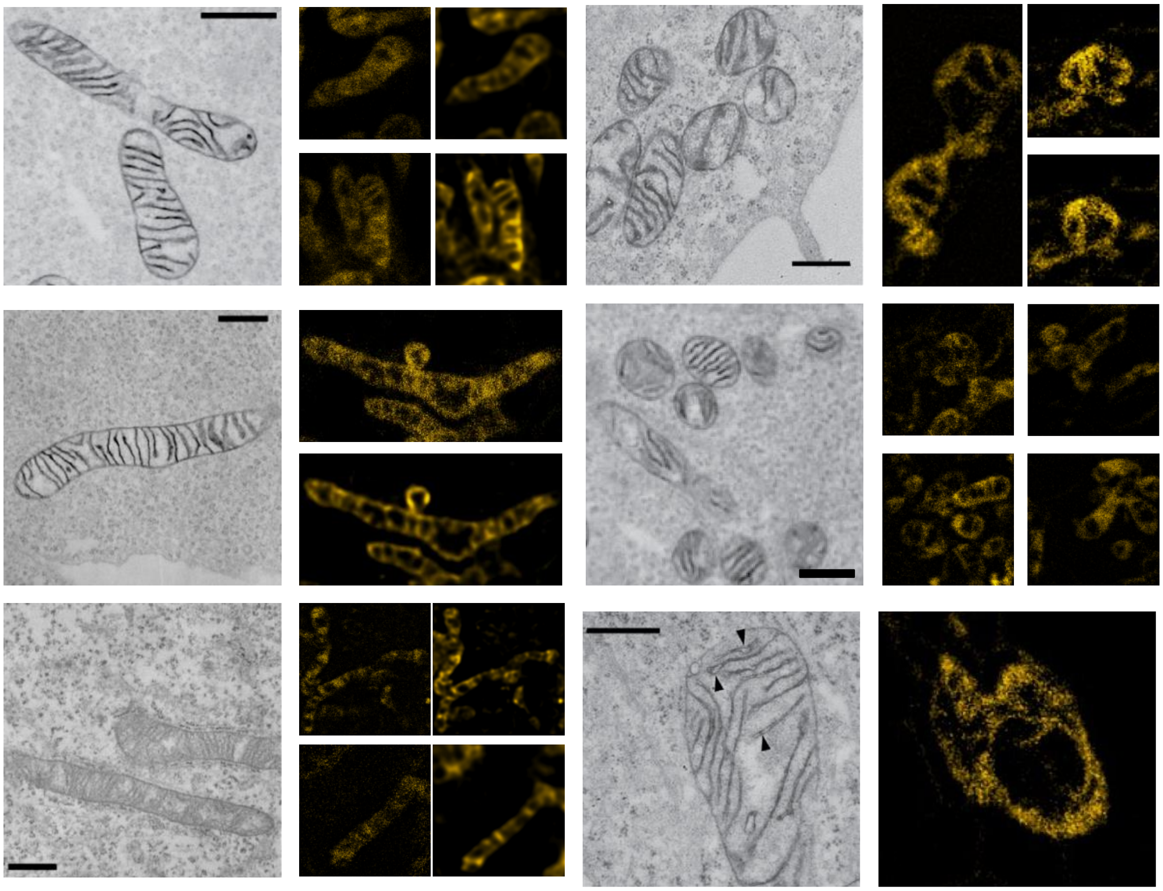

**Supplementary Figure 9.**

The comparison of ultra-fine structure of mitochondria were resolved by STED and EM[8,9]

[8] Lam S S, Martell J D, Kamer K J, et al. Directed evolution of APEX2 for electron microscopy and proximity labeling[J]. Nature methods, 12(1), 2015: 51.

[9] Martell, Jeffrey D., et al. "Engineered ascorbate peroxidase as a genetically encoded reporter for electron microscopy." *Nature biotechnology* **30**(11), *2012*: 1143.
