## Supplemental Notes for "Mitochondrial dynamics quantitatively revealed by STED nanoscopy with an enhanced squaraine variant probe"

### Supplementary notes

#### Supplementary note 1

##### Synthesis procedures of compounds

**Sample Preparation.** All reagents including solvents and chemicals used were reagent grade. All reactions were performed under the argon atmosphere with dry, freshly distilled solvents under anhydrous conditions. Silica gel (100-200 mesh) was used for flash column chromatography for purifications. Water used in all experiments was doubly purified by Milli-Q System equipment (Double De-ioned-water). The solutions of compounds were typically prepared from 1.0 mM stock solutions in DMSO.

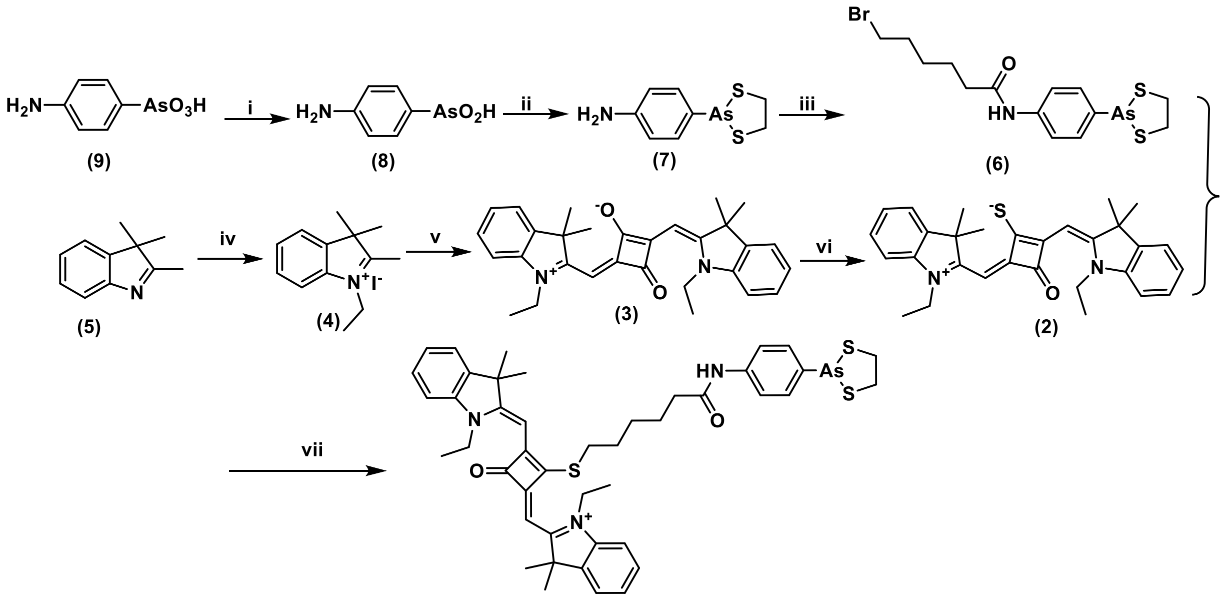


**Supplementary Scheme 1**. Synthetic route of target compounds of squaraine derivative (MitoESq-635); i) phenylhydrazine, ethanol, reflux for 2h, yield 78%; ii) ethylenedithiol, ethanol, reflux for 0.5 h, yield 47%; iii) 6-bromohexanoic, HATU, DMAP, DIPEA, dry DMF, room temperature, stirred for overnight, yield 60%; iv) ethyl iodide, reflux 12 hs in toluene, yield 79.6%; v) squaric acid, triethyl orthoforamate, reflux for 8 hs in ethanol, yield 65%; vi) Lawesson's reagent, stirred for 10 hs in dry DCM/THF at 40 $^oC$, 40%; vii) (**6**), NaI, stirred at 50 $^oC$ for 3 hs in anhydrous acetonitrile, 46%.

**[1] Synthesis of 4-aminophenylarsenoxide (8)**

Compound **8** was prepared according to the literature procedure. 4-aminophenylarsanilic acid **(9)** (11 g, 50 mmol) was dissolved in methanol (45 mL) and heated to reflux. Phenylhydrazine (12 mL, 110 mmol) was titrated dropwise to above mixture in 30 minutes. When $N_2$-production ceases, refluxing was continued for 2 hours. The mixture was condensed at 80 $^{o}$C to exclude the solvent, then washed with water (50 mL) and an aqueous NaOH solution (0.2 M, 30 mL) and finally with diethyl ether (2 × 80 mL). The aqueous solution was treated with aqueous $NH_4Cl$ solution (5 M, 40 mL) suspended overnight at 0 $^{o}$C. Precipitates were collected through a Büchner funnel filter, washed by ice-water (20 mL× 2) and dried over KOH to give **8** as white powder (9.5 g, 78% yield).

**[2] Synthesis of 2-p-aminophenyl-1, 3, 2-dithiarsenolane (7)**

To a solution of **8** (2.6 g, 13 mmol) in dry ethanol (25 mL), ethanedithiol (1.5 mL, 20 mmol) was added dropwise and heated to reflux for 30 minutes with constant stirring. The solution was chilled in dry ice/acetone, co-distilled with toluene (106 mL) and concentrated. A white crystal was recrystallized from ethanol yielding 2.5 g (47% yields) of **9**. $^{1}$H NMR (DMSO-$d_6$, 400 MHz): $\delta$ 3.13-3.36 (m, 4H), 5.40 (s, N-H, 2H), 6.56 (d, 2H, $J$ = 8.8 Hz), 7.27 (d, 2H, $J$ = 4.8 Hz) ppm.

**[3] Synthesis of 4-(6-bromohexanoylamidophenyl)-1, 3, 2-dithiarsenolane (6)**

In a 50 ml round bottom flask, 6-bromohexanoic acid (1.95 g, 10 mmol), HATU (2.5 g, 7 mmol), DMAP (0.12 g, 1 mmol) and DIPEA (0.5 mL) were added in dry DMF (15 mL) and the mixture was stirred for few minutes at room temperature. Then, 4-aminophenylarsenic **7** (1.3 g, 5 mmol) dissolved in DMF was added dropwise to above solution and stirring was continue overnight at room temperature. The reaction mixture was poured into saline solution (70 mL) and extracted with DCM (100 mL), dried over anhydrous $Na_2SO_4$. After evaporation of DCM, the product was purified by column chromatography to provide product **6** (2.6 g, yield 60%). $^{1}$H NMR ($CDCl_3$, 400 MHz): $\delta$ 1.45 (m, $CH_2$, 2H, $J$ = 8.0 Hz), 1.72 (m, $CH_2$, 2H, $J$ = 8.0 Hz); 1.85 (m, $CH_2$, 2H, $J$ = 8.0 Hz); 2.37 (t, $CH_2$, 2H, $J$ = 8.0 Hz); 3.14 (m, $CH_2$, 2H, $J$ = 4.0 Hz); 3.33 (t, $CH_2$, 2H, $J$ = 8.0 Hz); 3.38 (t, $CH_2$, 2H, $J$ = 8.0 Hz); 7.53 (dd, ArH, 4H, $J$ = 8.0 Hz); 8.06 (s, NH, 1H); $^{13}$C NMR ($CDCl_3$, 100 MHz): $\delta$ 24.87, 27.93, 32.61, 37.51, 42.06, 119.95, 131.69, 138.80, 139.13, 171.79 ppm.

**[4] N-ethyl-2, 3, 3-trimethylindolinium iodide (9)**

2, 3, 3-trimethyl-*3H*-indolenine (25 mL, 24.8 g, 156 mmol) and iodoethane (30 g, 192 mmol) were mixed in 50 mL dry toluene in 250 mL round flask, then the mixture was refluxed under argon atmosphere for 12 h, then stopped heating and cooled down. Then the precipitate was filtered through a Buchner funnel, the solid product was washed by diethyl ether and dried in vacuum to afford pink product (39 g, yield: 79.6%).

**[5] <u>Synthesis of ethyl-squaraine dye (3)</u>**

Squaric acid (1.14 g, 10 mmol) was placed in a 250 mL round bottom flask equipped with a suitable magnetic stirring bar, then triethylorthoformate (2 mL) and dry ethanol (100 mL) were added into the flask; the obtained reaction mixture was heated to reflux under argon atmosphere and stirred till the white squaric acid completely dissolved and the solution became transparent; and then N-ethyl-2, 3, 3-trimethylindolinium iodide (**9**) (7g, 22 mmol) was added into the reaction mixture, continued to react under refluxing, the reaction process was monitored by TLC, the reaction was quenched until all the starting materials was consumed to give deep blue solution. The reaction was cooled down and solvent was removed under vacuum evaporation to give raw product in dark-blue solid. The squaraine dye (3) was purified by silica column chromatography with the gradient eluent of DCM/n-hexane in blue solid (**3**) (2.9 g, 65%). $^{1}$H NMR (400 MHz, $CDCl_3$) $\delta$ 1.34 (t, $CH_3$, 6H, $J$ = 6.0 Hz); 1.73 (s, $CH_3$, 12H); 4.04 (q, $CH_2$, 4H, $J$ = 6.0 Hz); 5.92 (s, CH, 2H); 6.94 (d, ArH, 2H, $J$ = 8.0 Hz); 7.08 (t, ArH, 2H, $J$ = 8.0 Hz); 7.25 (t, ArH, 2H, $J$ = 8.0 Hz); 7.30 (d, ArH, 2H, $J$ = 8.0 Hz); $^{13}$C NMR (100 MHz, $CDCl_3$) $\delta$ 21.91, 27.18, 38.52, 49.53, 85.92, 109.59, 122.39, 123.91, 124.31, 128.41, 142.43, 169.78, 179.51, 181.02, 182.26.

**[6] <u>Synthesis of sulfido-ethyl-squaraine dye (2)</u>**

The above prepared blue solid (**3**) (1.3 g, 3 mmol) was placed in a round bottom flask (150 mL) dissolved in dry THF/DCM and Lawesson's reagent (2.2 g, 5 mmol) was added into above solution to provide greenish reaction, which was stirred at 40 $^{o}$C for 5 hs and monitored with TLC. The green spot was found to be on top of TLC plate with smaller polarity than dye (**3**). The reaction was cooled down to the room temperature. And then the solvent was removed by vacuum evaporation to give the residue with pungent smell, and the washed with saline water and the product was isolated *via* silica column chromatography to give sticky solid of (**2**) (0.56 g, 40%). $^{1}$H NMR (400 MHz, $CDCl_3$) $\delta$ 1.25 (s, $CH_3$, 6H); 1.38 (t, $CH_3$, 6H, $J$ = 6.0 Hz); 1.79 (s, $CH_3$, 6H); 4.05 (q, $CH_2$, 4H, $J$ = 6.0 Hz); 5.95 (s, CH, 2H); 7.00 (d, ArH, 2H, $J$ = 8.0 Hz); 7.15 (t, ArH, 2H, $J$ = 8.0 Hz); 7.35 (t, ArH, 2H, $J$ = 8.0 Hz); 7.58 (d, ArH, 2H, $J$ = 8.0 Hz).

**[7] <u>Synthesis of target compounds-As (MitoESq-635)</u>**

Sulfido-squaraine dye (2) (0.49 g, 1 mmol) was added into a round bottom flask (50 mL) in 20 mL dry acetonitrile, and ethylenedithiol-4-(6-bromohexanoylamido) phenylarsenicate (**6**) (0.18 g, 1.1 mmol) was added into above solution, which was heated at 50 $^{o}$C for 3 hs under the protection of argon atmosphere monitored with TLC. Till complete disappearing of the starting materials, the reaction was cooled down to room temperature and the solvent was evaporated in vacuum. The residue was separated through silica column chromatography with the eluent of DCM/ methanol to give the blue solid (**1**) (0.38 g, 46% yield). $^{1}$H NMR (400 MHz, $CDCl_3$) $\delta^{1}$H NMR ($CDCl_3$, 400 MHz): $\delta$ 1.43 (t, $CH_3$, 6H, $J$ = 8.0 Hz), 1.70 (s, $CH_3$, 12H); 1.83 (m, $CH_2$, 2H, $J$ = 6.0 Hz); 2.10 (m, $CH_2$, 2H, $J$ = 6.0 Hz); 2.72 (t, $CH_2$, 2H, $J$ = 6.0 Hz); 3.11 (m, $CH_2$, 2H, $J$ = 4.0 Hz); 3.28 (m, $CH_2$, 2H, $J$ = 4.0 Hz); 3.63 (t, $CH_2$, 2H, $J$ = 6.0 Hz); 4.21 (q, $CH_2$, 4H, $J$ = 8.0 Hz); 5.72 (s, CH, 2H); 7.17 (d, ArH, 2H, $J$ = 8.0 Hz); 7.27 (t, ArH, 2H, $J$ = 8.0 Hz); 7.37 (d, ArH, 2H, $J$ = 8.0 Hz); 7.46 (d, ArH, 2H, $J$ = 8.0 Hz); 7.96 (d, ArH, 2H, $J$ = 8.0 Hz); 9.69 (s, NH, 1H); $^{13}$C NMR ($CDCl_3$, 100 MHz): $\delta$ 12.57, 25.19, 26.17, 27.59, 30.00, 32.27, 37.24, 40.74, 41.80, 50.64, 88.32, 111.49, 120.01, 122.58, 126.41, 128.87, 131.19, 137.11, 140.59, 141.01, 142.53, 171.82, 172.01, 173.08, 173.43, 174.15, 175.74 ppm; ESI-MS (m/z): $[M]^+$ calculated for $C_{44}H_{51}AsN_3O_2S_3^+$, 824.2; found, 824.0.

#### Supplementary note 2

##### MitoESq-635 covalently binding behavior to vicinal dithiols in mitochondria

As seen in Supplementary Figure 3, the probe was first incubated with commercial available VDP of thioredoxin, known as a kind of mitochondrial proteins containing a pair of dithiol residue in proximity on the surface of its reduced form, which can adjust mitochondrial redox homeostasis, under different conditions, the SDS-PAGE experimental results demonstrated that the probe can covalently attach to the protein in the presence of DTT. Moreover, MitoESq-635 can also bind to the proteins with vicinal dithiols in mitochondria of HeLa cells. The covalently binding behavior was verified *via* the comparison of fluorescence changes of MitoESq-635 in live HeLa cells, fixed cells and the washed fixed cells with pure ethanol, respectively (Supplementary Figure 4). Additionally, the colocalization experiment of MitoESq-635 with Rhodamine123 in HeLa cells were carried out to further identify the covalent binding of MitoESq-635 to mitochondrial VDPs As shown in Supplementary Figure 5, in the live HeLa cells, the probe indicated well fluorescent overlapping with that of Rhodamine 123; whereas, when the cells were fixed with glutaraldehyde upon incubation with both probes for 30 minutes, MitoESq-635 was still kept in mitochondria without marked diffusion, not like Rhodamine 123 diffusing all over the cells and fluorescence decreasing markedly, which again substantiate the covalent binding of MitoESq-635 to VDPs in mitochondria.

#### Supplementary note 3

##### Possibility for wide applications in fluorescent labeling of enhanced squaraine dyes derivatives

To verify the wide applications in cellular imaging of the newly developed enhanced squaraine dyes (ESq), the sulfide-squaraine dyes (**2** in Supplementary Scheme) were modified with different targeting ligand to carry out wide applications of STED imaging for biological targets in live cells. As demonstrated above, the alkyl sulfide-squaraine was suitable for STED imaging in live cells, as it was capable of STED imaging under low power of depletion laser and endurance of longtime laser illuminating, which revealed to be better than ATTO647N in STED imaging. The intracellular selective labeling of the probes mainly resorts to the targeting ligands, different targeting ligands conjugating with the sulfide-squaraine dye (**2**) will guide the obtained fluorescent probes to exactly mark the biological objects. As shown in the following figure, a new protocol was designed to prepare fluorescent probes for various objects. The intermediate sulfide-squaraine (**2)** was the first subject to reacting with haloalkyl acid to make a platform compound (**10)**, which can react with different targeting ligands with amines to provide target probes. The group R could be a lot of functional ligands for different objects inside cells, such as para-aminophenyl-arsenicate[1], phalloidine[2], glibenclamide[3], ceramide[4], indomethacin[5], and other antibodies or inhibitors *etc.*, to specifically label VDPs, cell backbone, ER, Golgi apparatus or single cyclooxygenase (COX) protein or other enzyme inside cells, respectively, which could achieve wide applications of the probes in live cell STED imaging strategies.

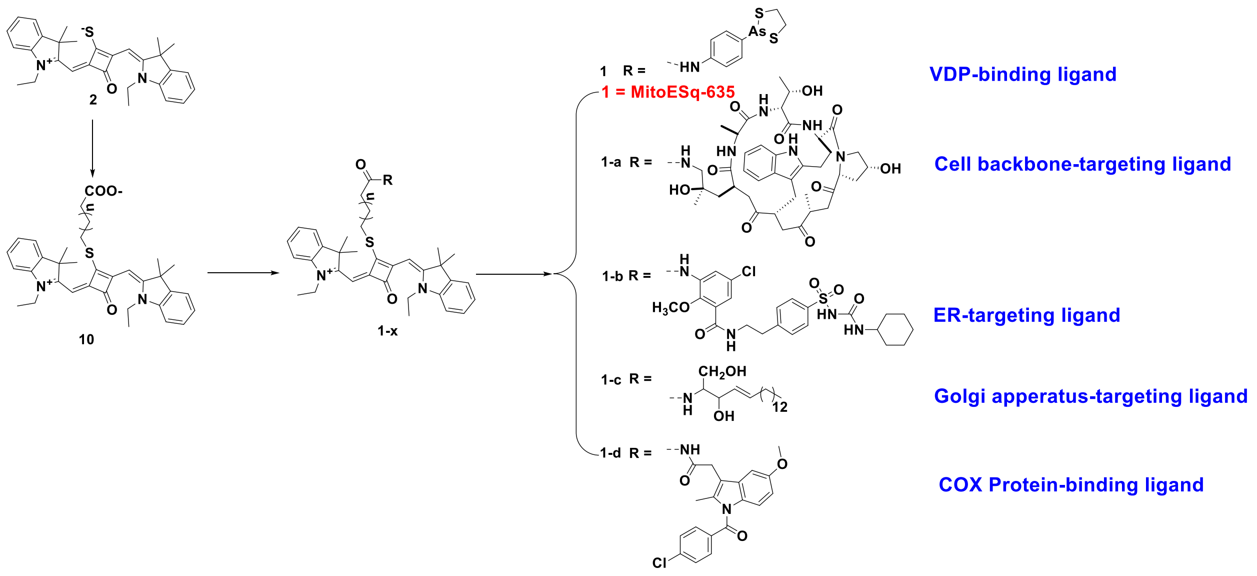


**Supplementary Notes Figure 1** Designing strategies for fluorescent probes for wide applications in STED imaging of live cells.

#### Supplementary note 4

##### Photo-toxicity in super-resolution microscopy

The real promise of super-resolution is the ability and the hope of living cell dynamics imaging. But it's of big challenges to fulfill this promise. In order to map subcellular structure at nanoscale, much more photon should be attained for get a decent signal to background ratio. Cells have a limited tolerance for light intensity because of photodamage and photo-toxicity. However, these super-resolution methods including (d) STORM/PALM and STED microscopy require high doses of light to achieve resolution enhancement breaking diffraction limit. Even though there have been lots of super resolution technical demonstrations in living cells, there's been little real biology learned.

Photo-toxicity depends on both light dose and wavelength, as well as the site environment and cell type of employed fluorophore. Generally, lower laser power and longer wavelength are decent strategies to avoid unwanted photo-toxicity. Also, the acquisition speed of these methods (fourth column) is far slower than the biological dynamics in living cells, so it is difficult to get motion-induced artifacts, which means no single parameter can be optimized without compromising the others to technical consideration. To get effective information by bioimaging, images with enough contrast, spatial, and temporal resolution should be attainable while leave the biology of interest intact. Contrast, spatial and temporal resolution are dependent t to each other. Usually, better contrast, higher spatial or temporal resolution requires much higher light (excitation or modulation) dose and/or more fluorescence photons, while intense light illumination will cause marked photo damage/toxicity in turn. Minimizing photo damage/ toxicity is the most key strategy to carry out living cell imaging because any imaging of biological dynamics/ process will come to the end once the photo-toxicity occurs in the samples. Therefore, the probe which can achieve enough contrast, spatial resolution, and temporal resolution with modest light dose (excitation or modulation) is highly desired. Considering the fluorescence photon budget for living cell imaging, luminescent probes should be excited under low power of light and have enough emission photons for a decent signal to noise ratio. MitoESq-635 has excellent photophysical property over golden standard organic dye ATTON 647 N probe for STED nanoscopic imaging because it has excitation and depletion spectrum in red and infrared as well as lower saturated intensity.

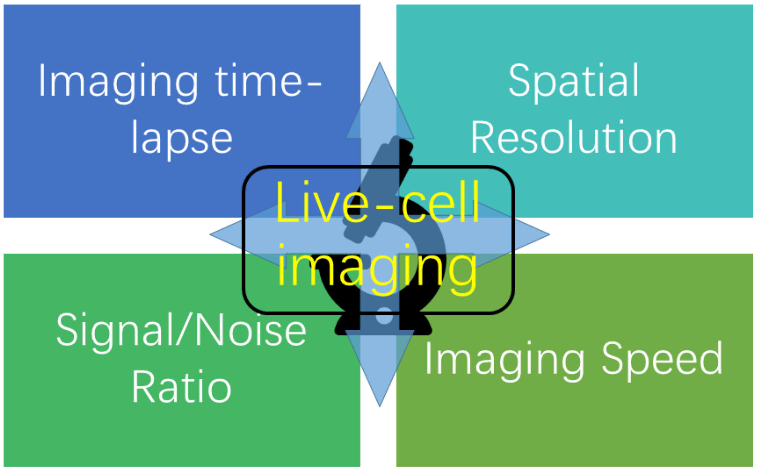


**Supplementary Notes Figure 2** The four main considerations for live imaging.

Photon-induced cell damage is thus a crucial consideration for light microscopy of living samples, and minimizing it is a fundamental concern. Many factors can influence phototoxicity including fluorophores (concentration and their subcellular localization), excitation wavelength and intensity (peak and time-averaged), exposure time (scanning time and the amount of dark recovery time between images), sample preparation, media, singlet oxygen concentration, cell type and age, the developmental stage of the organism, and synergistic effects of experimental perturbations, which can all affect a live sample under fluorescence microscopic observation.

It is a complex phenomenon consisting of wavelength-dependent photophysical mechanisms that can generate highly reactive photochemical products, heat, and DNA damage. The phototoxicity by the damaging radicals produced by laser excitation can be minimized, but unavoidable. Cells can tolerate laser illuminating as long as their defense mechanisms are not overwhelmed. During the fluorescent imaging, fluorescent proteins or dyes in their excited state are readily to generate reactive oxygen species (ROS). These unstable, short-lived reactive species may in turn damage the chemical structure and biological function of proximal biomolecules, so any type of fluorescence microscopy risks causing light-induced damage to living samples. The degree to which illumination causes photodamage depends strongly on the sample tolerance, the duration of the observation, and the observed cellular process.

Mitochondria are intricately involved in the activation and regulation of different pathways adjusting the balance between cell death and survivals. Complexes I and III in the inner mitochondrial membrane are the key producers of ROS that plays crucial roles in mitochondrial biology. Mitochondrial membranes become depolarized at excessive ROS and may be destroyed in a ROS burst by the prolonged opening of mitochondrial permeability transition pores. This mitochondrial permeability transition can further induce DNA fragmentation and cell apoptosis. The mitochondrial membrane contains key regulators (cytochrome c, Bcl-2 family of proteins), which play vital roles in the homeostasis between apoptosis and survival. Even at low irradiance, mitochondria-derived ROS may lead to cell death. One possible reason is that mitochondria are rich in chromophores that are photosensitive in the UV and visible wavelength range, such as heme proteins, flavoproteins, and NADH or NADPH. This is compounded by the complex interaction of mitochondria with the ER. Such interactions leading to cell death are exploited in photodynamic therapy, where selective photosensitizers are used to generate large amounts of ROS through illumination. This field is also a rich source for studies of localized ROS damage to cell populations, tissues, and organisms. Although specimens show diverse signs of light-induced damage, common themes do exist. A frequent recommendation for assessing phototoxicity is to look for telltale morphological signs such as cellular swelling and rounding, blebbing, or the appearance of vacuoles.

#### (III) Chemical Structure Identifications

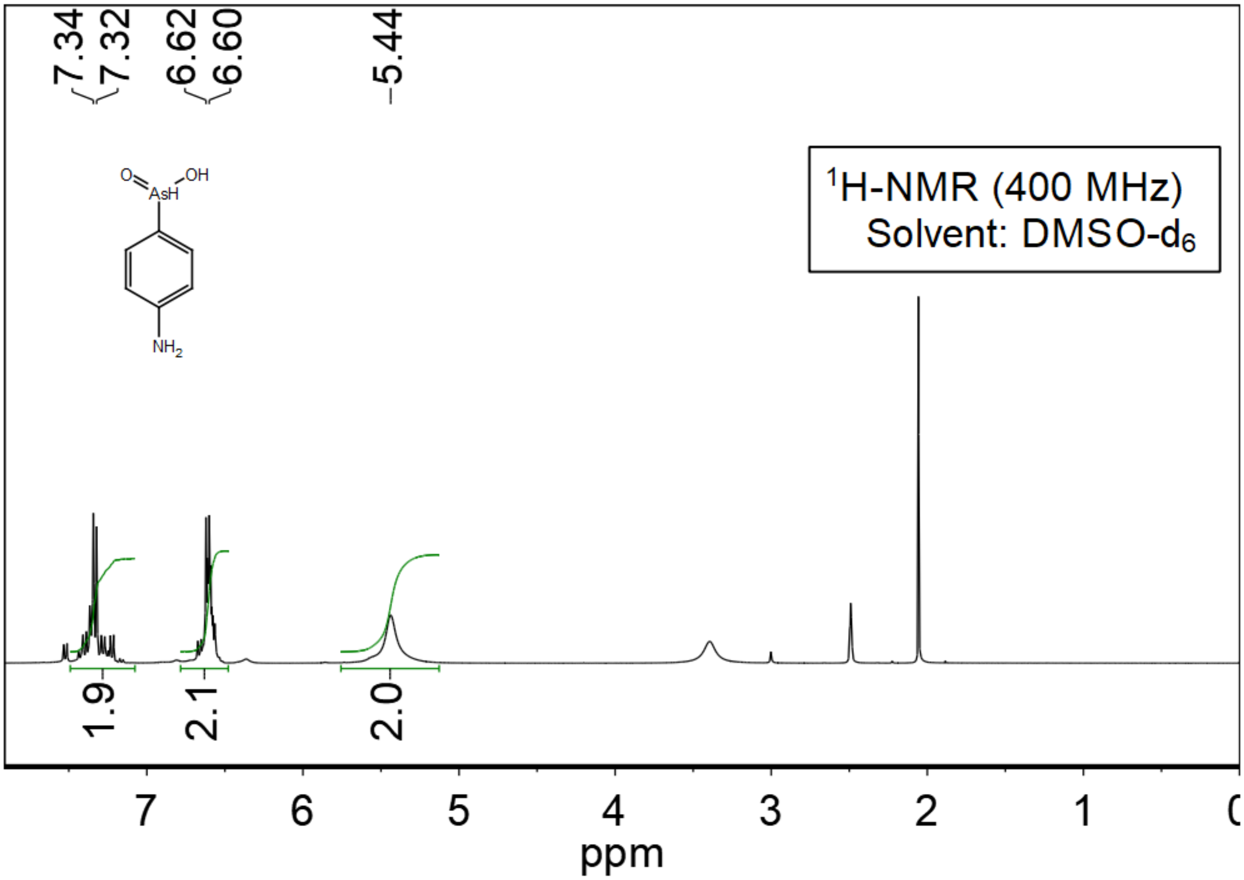


**Supplementary Notes Figure 3**

Chemical Structure Identifications of compounds

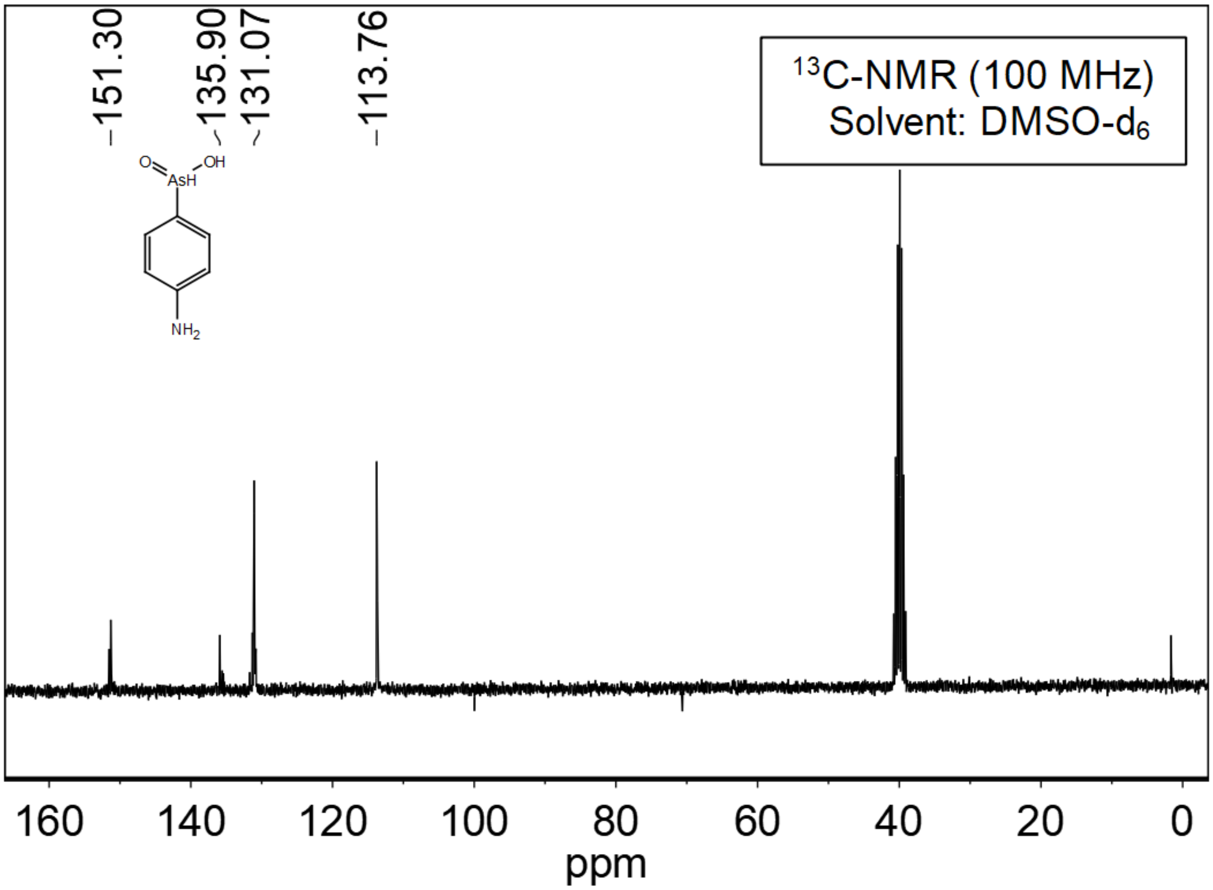


**Supplementary Notes Figure 4**

$^{1}$H and $^{13}$C NMR spectra of compound 8

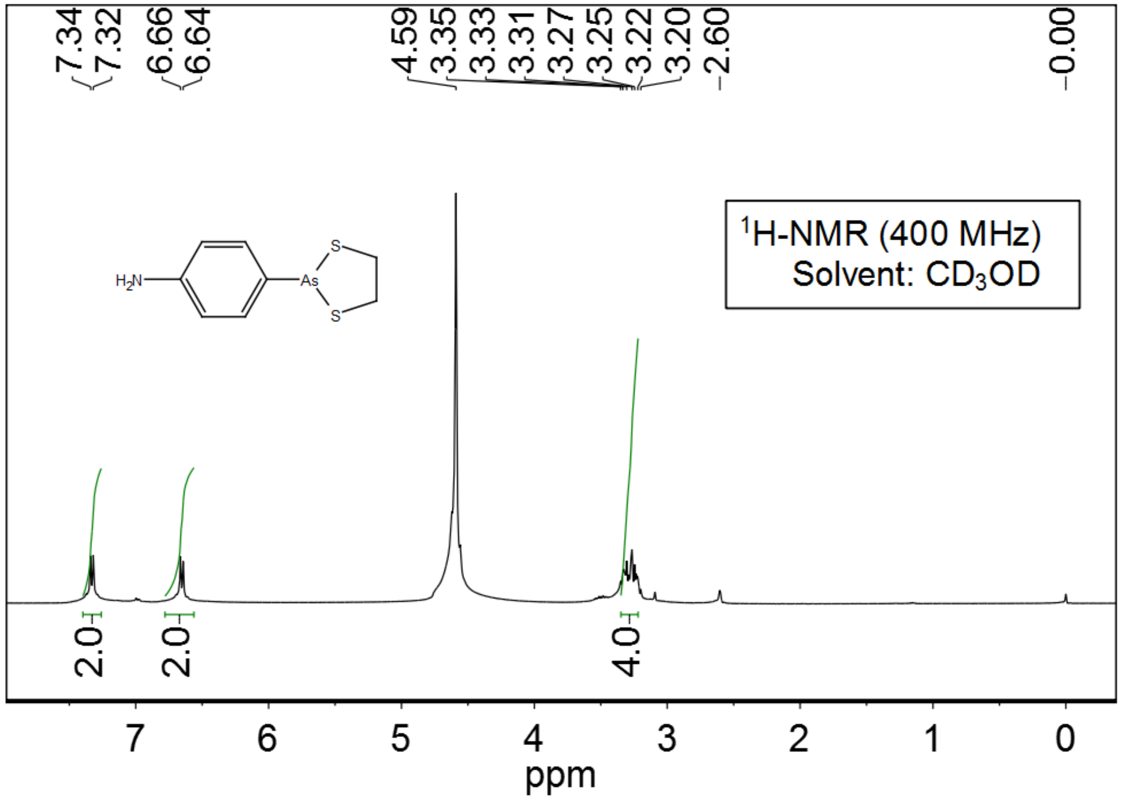


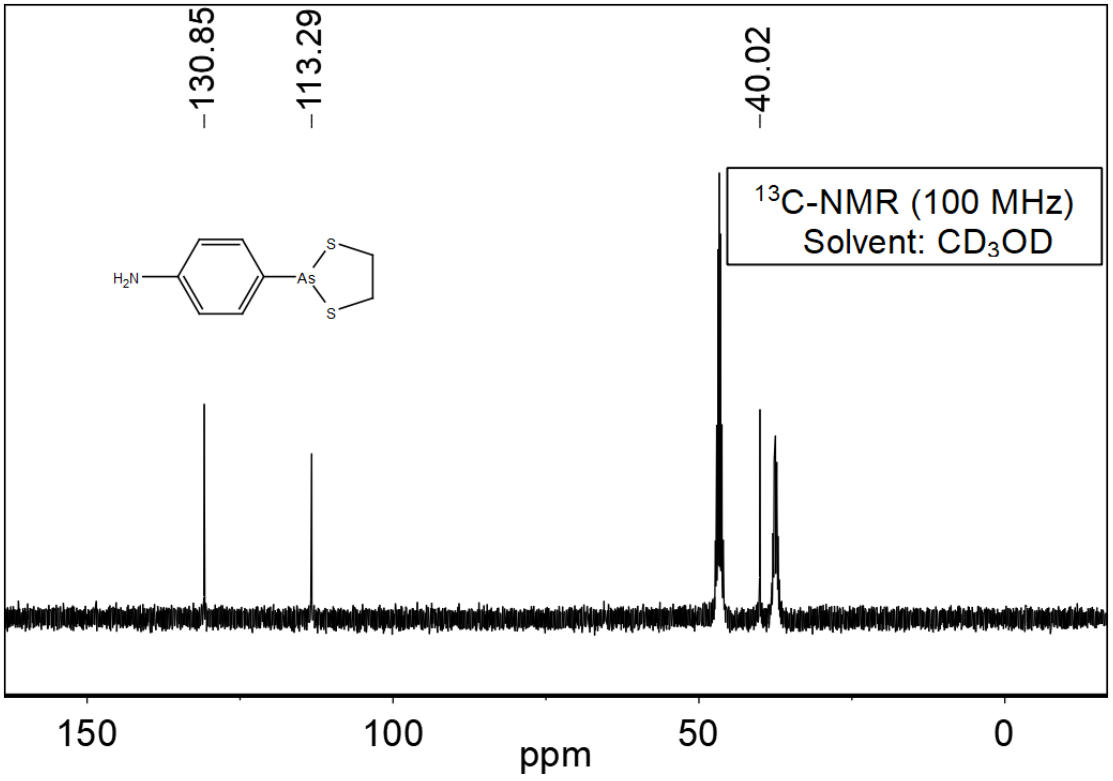


**Supplementary Notes Figure 5**
$^{1}$H and $^{13}$C NMR spectra of compound 7

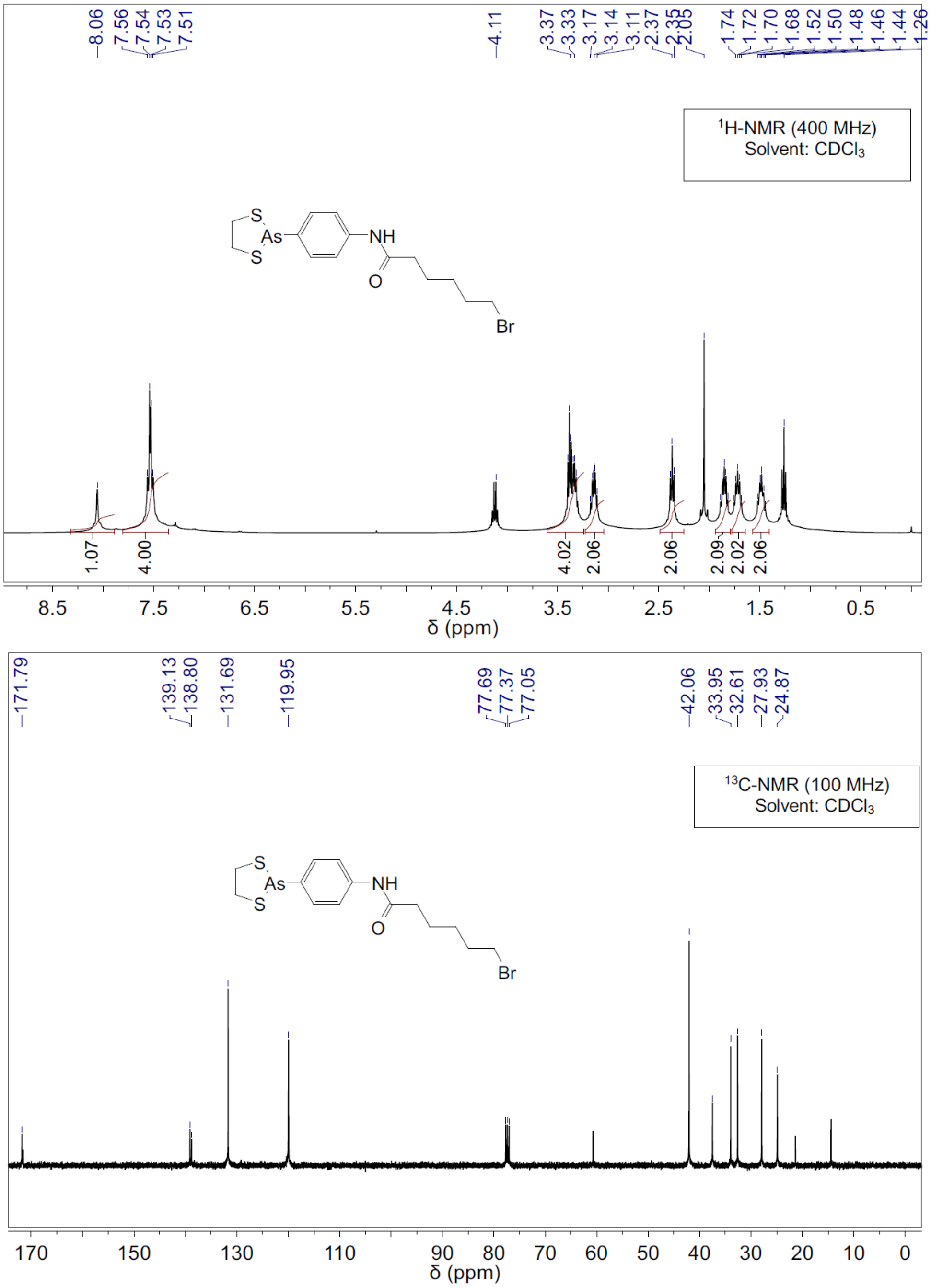


**Supplementary Notes Figure 6**

$^{1}$H and $^{13}$C NMR spectra of compound 6

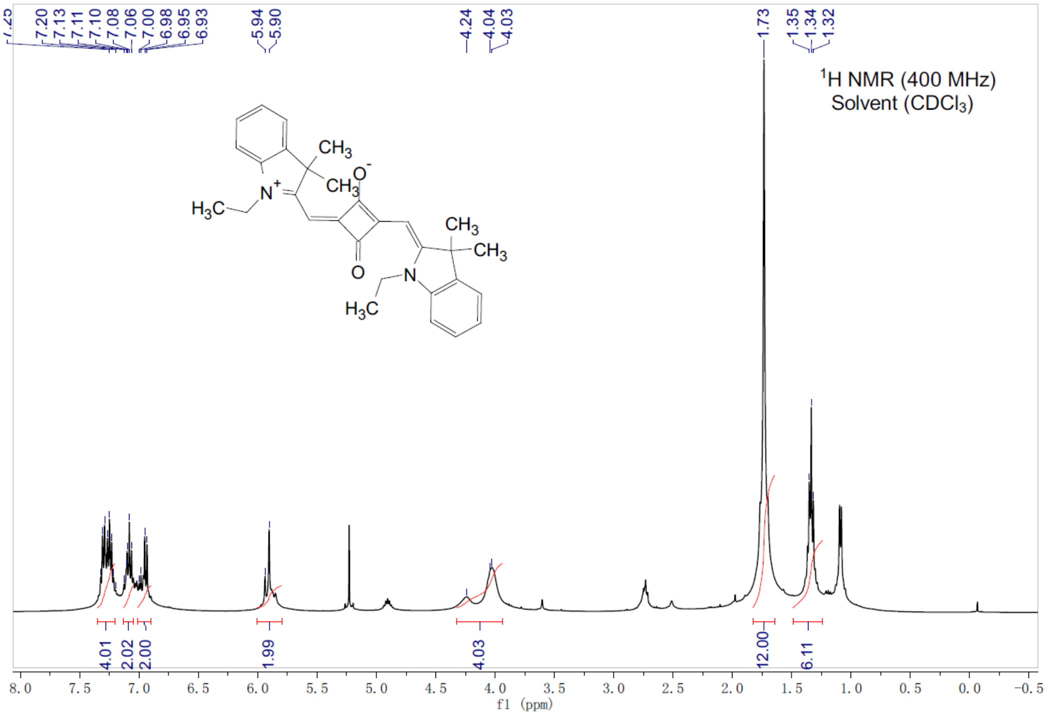


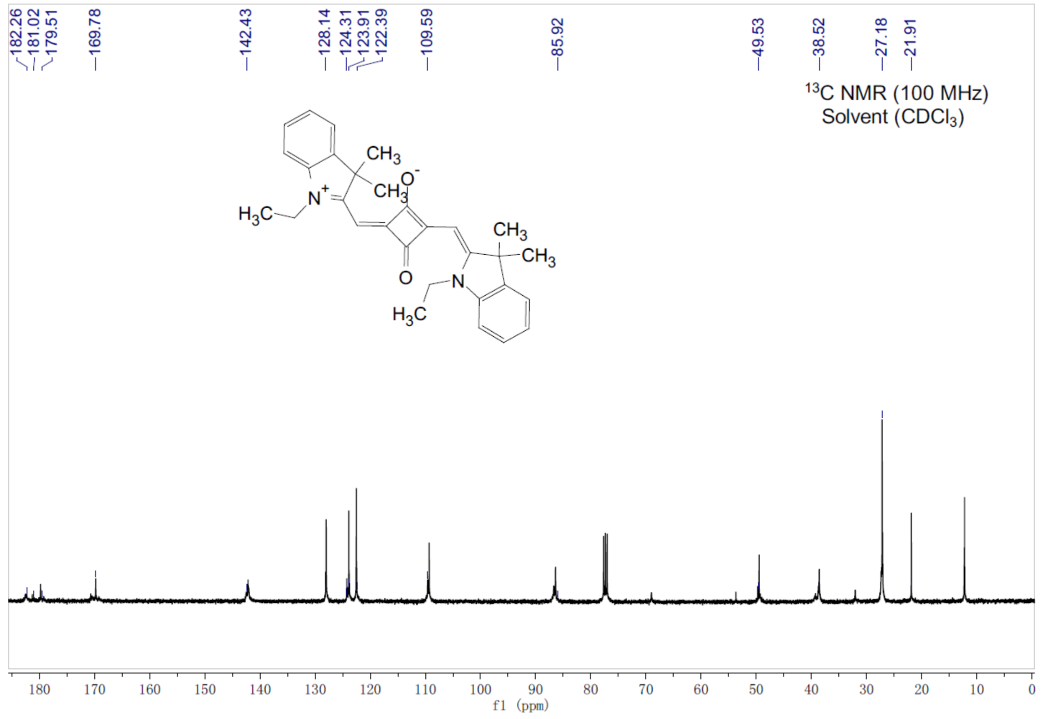


**Supplementary Notes Figure 7**

$^{1}$H and $^{13}$C NMR spectra of compound 3

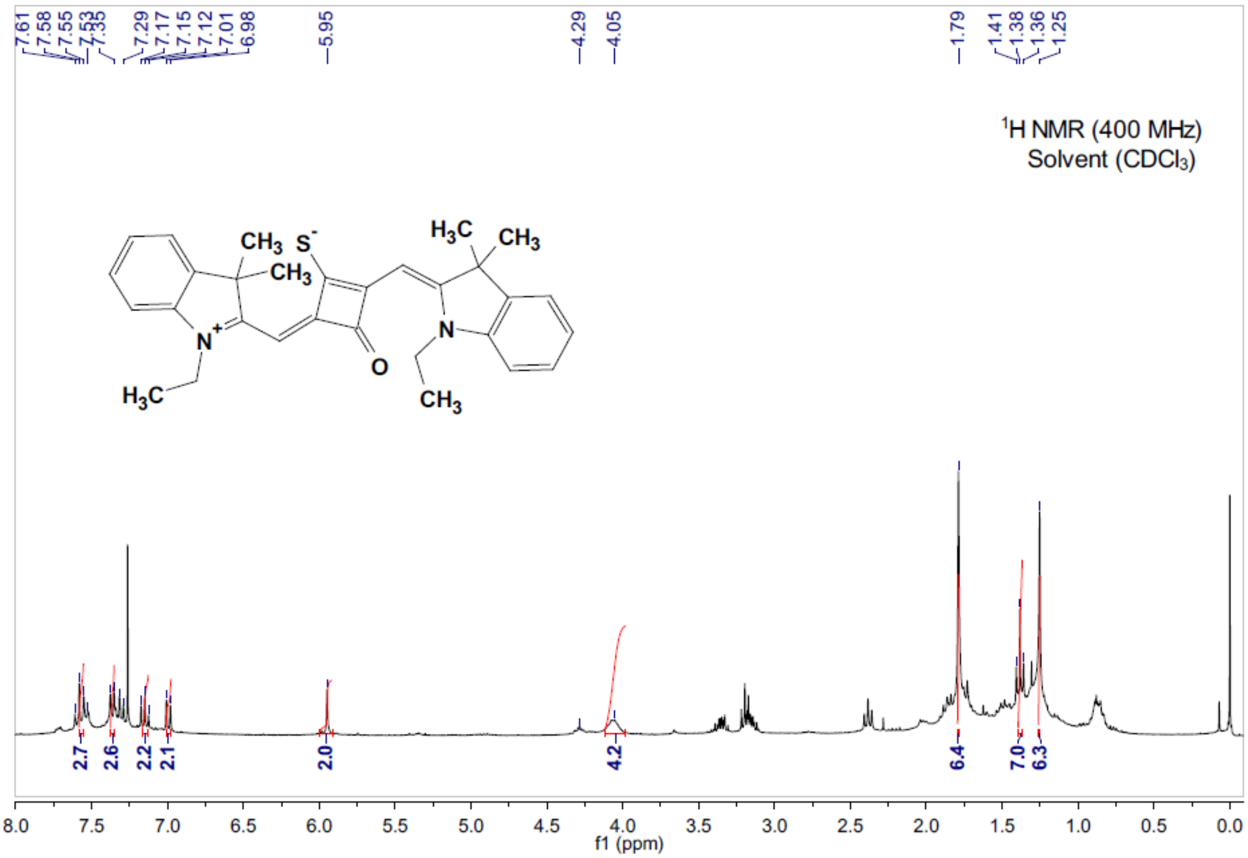


**Supplementary Notes Figure 8**

$^{1}$H spectra of compound 2

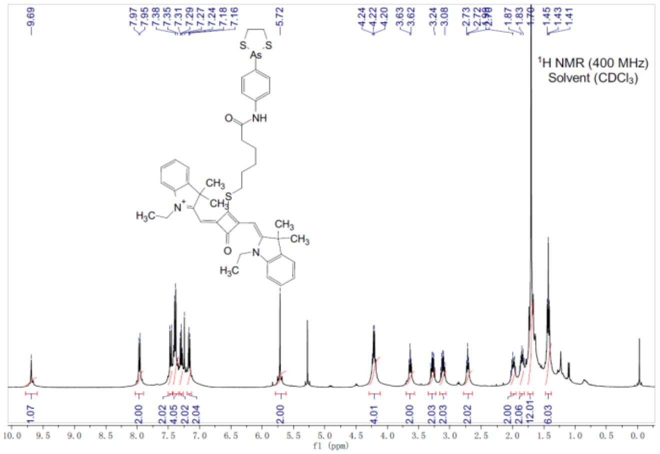


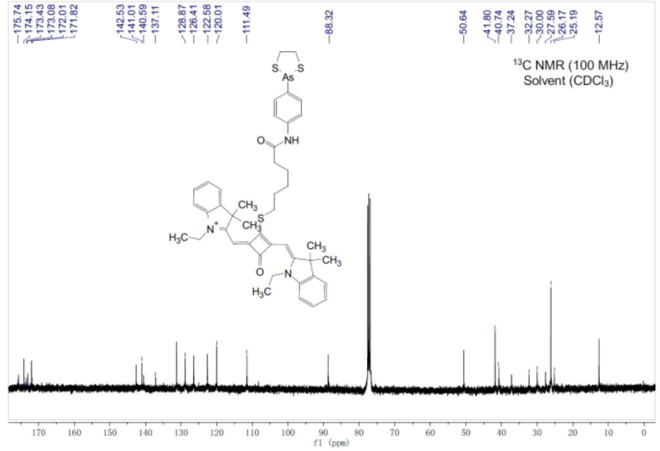


==== Shimadzu LabSolutions Data Report ====

<Spectrum>

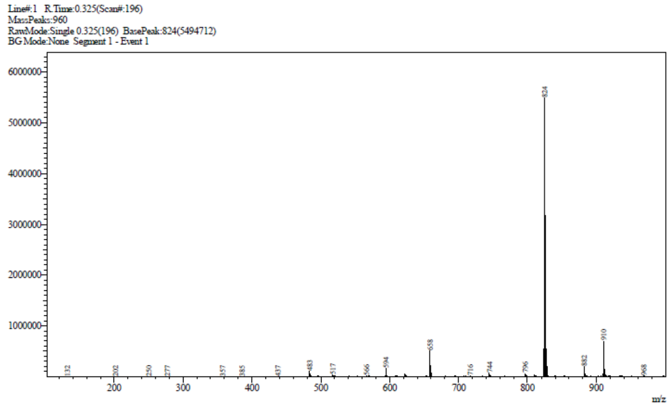


**Supplementary Notes Figure 9**

$^1$H, $^{13}$C NMR and mass spectra of compound
